# Structure and mechanism of LARGE1 matriglycan polymerase

**DOI:** 10.1101/2022.05.12.491222

**Authors:** Soumya Joseph, Nicholas J. Schnicker, Zhen Xu, Tiandi Yang, Jesse Hopkins, Maxwell Watkins, Srinivas Chakravarthy, Omar Davulcu, Mary E. Anderson, David Venzke, Kevin P. Campbell

## Abstract

Matriglycan is a linear polysaccharide of alternating xylose and glucuronate that binds extracellular matrix proteins and acts as a receptor for Lassa fever virus. LARGE1 synthesizes matriglycan on dystroglycan and mutations in *LARGE1* cause muscular dystrophy with abnormal brain development. However, the mechanism of matriglycan polymerization by LARGE1 is unknown. Here, we report the cryo-EM structure of LARGE1. We show that LARGE1 functions as a dimer to polymerize matriglycan by alternating activities between the xylose transferase domain on one protomer and the glucuronate transferase domain on the other protomer. Biochemical analyses using a recombinant Golgi form of dystroglycan reveal that LARGE1 polymerizes matriglycan processively. Our results provide mechanistic insights into LARGE1 function and may facilitate novel therapeutic strategies for treating neuromuscular disorders or arenaviral infections.

**One-Sentence Summary:** Dimeric LARGE1 processively polymerizes matriglycan on dystroglycan using orthogonal active sites on alternate protomers.

## Introduction

Matriglycan is a linear polysaccharide of alternating xylose and glucuronate that is polymerized on dystroglycan (*1*), which is a mucin-like protein that is part of the dystrophin-glycoprotein complex (DGC) (*2*). Matriglycan binds laminin-globular (LG) domain-containing proteins in the extracellular matrix (*3,4*) to link the cytoskeleton with the basement membrane via the DGC (*2, 5-6*). The absence or reduction of matriglycan can cause various forms of muscular dystrophy with or without abnormalities in brain development (*7, 8*). Additionally, matriglycan is a receptor for Lassa fever virus (LASV) and other Old-World mammarenaviruses (*9, 10*). Matriglycan polymer length appears to be cell-type specific and is physiologically regulated during development and regeneration (*11*) but can be pathologically decreased by the expression of LASV or the related lymphocytic choriomeningitis virus spike glycoproteins (*12*).

The bifunctional glycosyltransferases, LARGE1 and LARGE2 transfer alternating xylose and glucuronate monosaccharides repetitively to polymerize matriglycan on human dystroglycan (DAG1) residues threonine-317, 319 and 379 (*11, 13, 14*), that carry a phosphorylated core M3 *O*-glycan stem (*15*). Mutations in *LARGE1* are rare but can cause congenital muscular dystrophies (*16*) with severe intellectual disability (*17*). Interestingly, GWAS studies found single nucleotide polymorphisms (SNPs) in *LARGE1* are common (35%) in the West African Yoruba population of Nigeria where LASV is endemic, but absent among the Luo and Masai people of Kenya and Tanzania where LASV is not found, suggesting that the *LARGE1* gene has undergone selection in populations where LASV is endemic (*18*).

LARGE1 is a single-pass type II transmembrane protein localized to the Golgi apparatus (*19*). The lumenal portion consists of a putative coiled-coil domain and two globular catalytic domains. The tandem, but independent, glycosyltransferase-A (GT-A) domains, adopt a Rossmann-like fold, that typically coordinate a manganese co-factor in each active site using signature double-aspartate sidechains (DXD) and a C-terminal histidine sidechain (*20*). Despite knowing the domain architecture, it is unclear how LARGE1 polymerizes matriglycan on dystroglycan. We show that LARGE1 proteins form dimers and that LARGE1 polymerizes matriglycan, of defined length, processively on dystroglycan by using the xylose transfer activity on one protomer with the glucuronate transferase on the other. Determining the structure of LARGE1 and the mechanism of matriglycan polymerization on dystroglycan may form the basis for managing matriglycan-deficient neuromuscular disease and LASV infections.

## Results

### LARGE1 is a glycosylated dimer

We performed light scattering analysis using multi-angle light scattering (MALS) and small-angle X-ray scattering (SAXS) to determine the oligomeric state of LARGE proteins and to obtain low-resolution molecular envelopes. *LARGE2*, which is a gene duplication product of *LARGE1*, does not encode the coiled-coil domain (*19, 20)*. We analyzed LARGE1ΔTM and LARGE2ΔTM in parallel to determine whether the coiled-coil domain was responsible for multimerization (Fig. 1A). Light scattering analysis showed that soluble recombinant constructs of LARGE proteins, in which the transmembrane domain was replaced by 3xFLAG (*13*), form dimers in solution (Fig. 1A–C and Table 1). LARGE2ΔTM also forms a dimer, despite the absence of a coiled-coil domain (Fig 1A and S1), suggesting that the globular catalytic domains of LARGE proteins are sufficient for self-association. The corrected Porod volume from SAXS data are also consistent with LARGE proteins forming dimers. SAXS-derived molecular envelopes show that LARGE1ΔTM dimers are parallel and consist of a globular head and a stem that is presumably the density representing the coiled-coil domain (Fig. 1C). The maximum dimension of LARGE1ΔTM is ~180 Å calculated from the pair-distance distribution function (Table 1; D_max_ and Fig. S2; P(r)). The parallel orientation of LARGE dimers is consistent with their biological topology as type-II single-pass transmembrane proteins in which the globular catalytic heads protrude into the Golgi lumen (Fig. 1D).

**Fig. 1.**
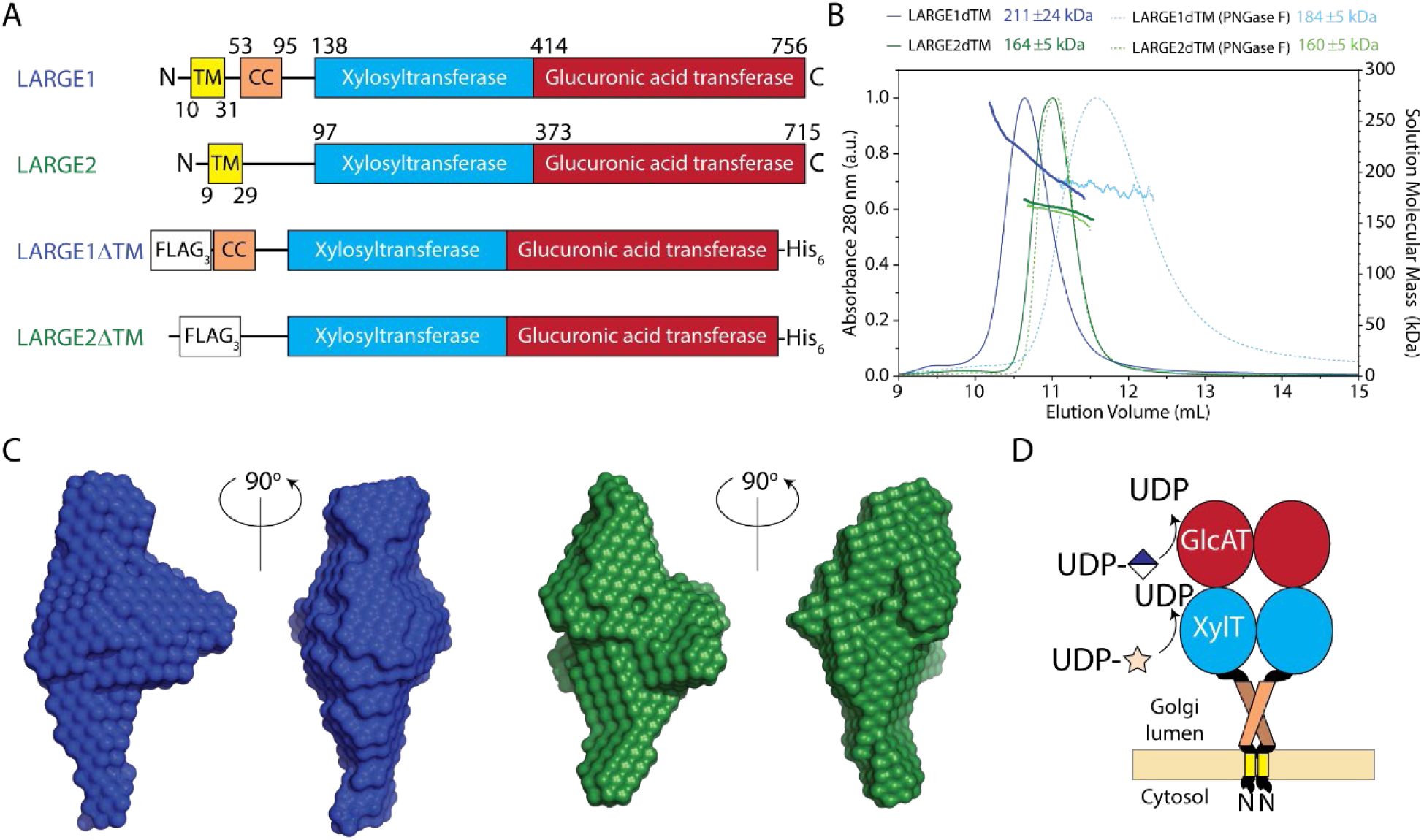
Light scattering analyses of LARGE1ΔTM and LARGE2ΔTM. (**A**) Schematic of LARGE proteins. LARGE2 lacks a coiled-coil domain (CC) found in LARGE1. Transmembrane domains (TM) were replaced with 3xFLAG tag (FLAG_3_) and a C-terminal hexahistidine tag was added (His_6_). (**B**) Size-exclusion chromatogram (SEC) with on-line multi-angle light scattering (MALS) of LARGE1ΔTM (blue) and LARGE2ΔTM (green) or treated with PNGase F (dotted blue and light green, respectively). The elution profile was monitored using absorbance at 280 nm (continuous line) over the elution volume (mL). Discrete points represent solution weight-average molecular weight (MW, kDa) calculated using a change in refractive index *dn/dc* = 0.185 for protein and *dn/dc* = 0.135 for glycan. (**C**) Molecular envelopes of LARGE1ΔTM (blue) and LARGE2ΔTM (green) from small-angle X-ray scattering (SAXS). (**D**) Probable organization of dimeric LARGE1 in the Golgi membrane.

**Table 1.**
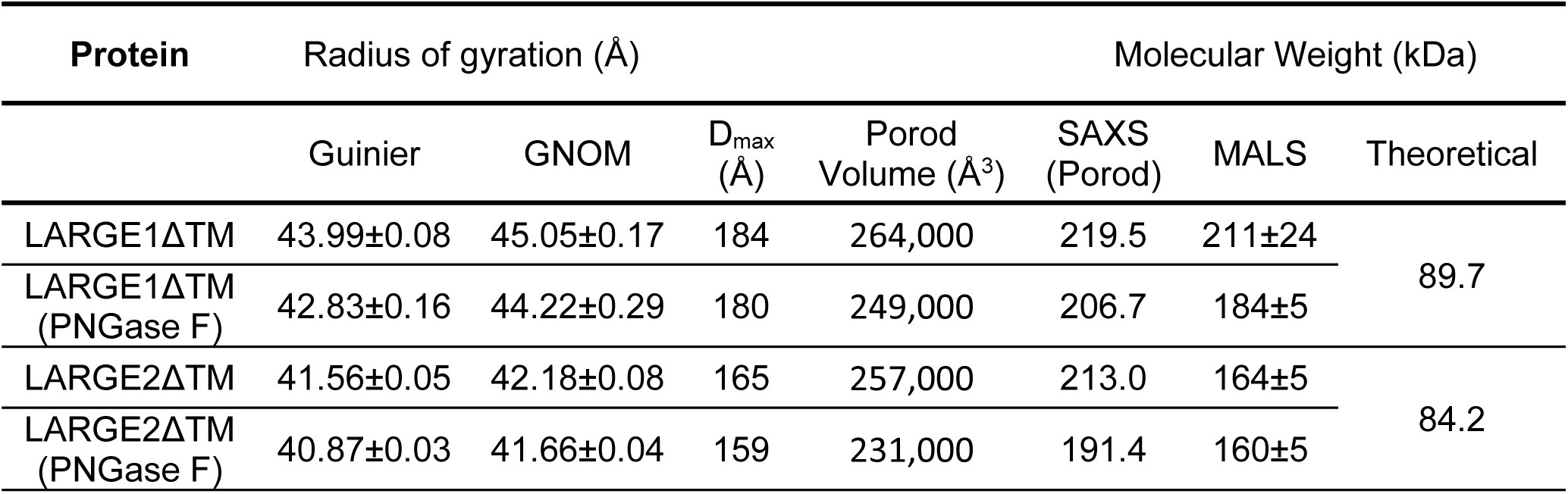
Biophysical parameters of LARGE proteins derived from multi-angle light scattering (MALS) and small-angle X-ray scattering (SAXS). The radius of gyration was estimated in reciprocal space from Guinier fitting and in real space from GNOM. The maximum dimension (D_max_) was estimated from the pair-distance distribution function (P(r)). The corrected Porod volume and molecular weight obtained from SAXS are calculated from Porod’s law. The solution weight-average molecular weight from MALS can be compared to the protein-only theoretical monomer weight.

Proteins in the secretory pathway are often glycosylated, which increases their molecular weight above the theoretical protein-only molecular weight that is predicted from the primary sequence. Biochemical and protein-conjugate analyses by MALS and mass spectrometry were used to determine the degree of glycosylation. The weight-average molecular weights of LARGE constructs were slightly higher than double the theoretical monomeric protein-only molecular weight (Table 1), suggesting that up to 20% of the observed molecular weight may be attributed to glycan. We observed an oligomeric state closer to an integer of two for LARGE proteins treated with PNGase F suggesting that N-glycan modifications contribute to the non-integral oligomeric state (Fig. 1B, Table 1 and Fig. S3) observed in untreated samples. A decrease in molecular weight across the elution peak for untreated LARGE protein can be partially attributed to heterogeneous glycosylation. Although the size of LARGE proteins decreases when N-glycans are removed, dimerization is unaffected (Fig. 1, Table 1).

### Cryo-EM reconstruction of LARGE1

A volume representing globular catalytic domains of LARGE1ΔTM was reconstructed using single-particle cryo-EM (Fig. 2 and Fig. S5). The cryo-EM reconstruction confirms that LARGE1ΔTM is a dimer. Each protomer consists of tandem Rossmann folds, which have a core composed of seven β-sheets surrounded by α-helices – a classic αβα-sandwich. Additionally, conservation of surface residues on each LARGE protomer primarily maps to the dimer interface (Fig. S4B). Although we reconstructed LARGE1ΔTM in both C1 and C2 symmetries, the homodimer is not perfectly symmetric. One xylose transferase domain in each dimer remained incompletely resolved, presumably due to a high degree of flexibility relative to the remainder of the protein. Approximately 16% of chain A and 8% of chain B are outside the map density. The coiled-coil domain and N-terminal tail were not resolved likely because they were averaged to the level of noise during particle alignment due to their dynamics with respect to the globular catalytic head. For similar reasons, the N-glycans on LARGE1ΔTM were not resolved in the cryo-EM reconstruction.

**Fig. 2.**
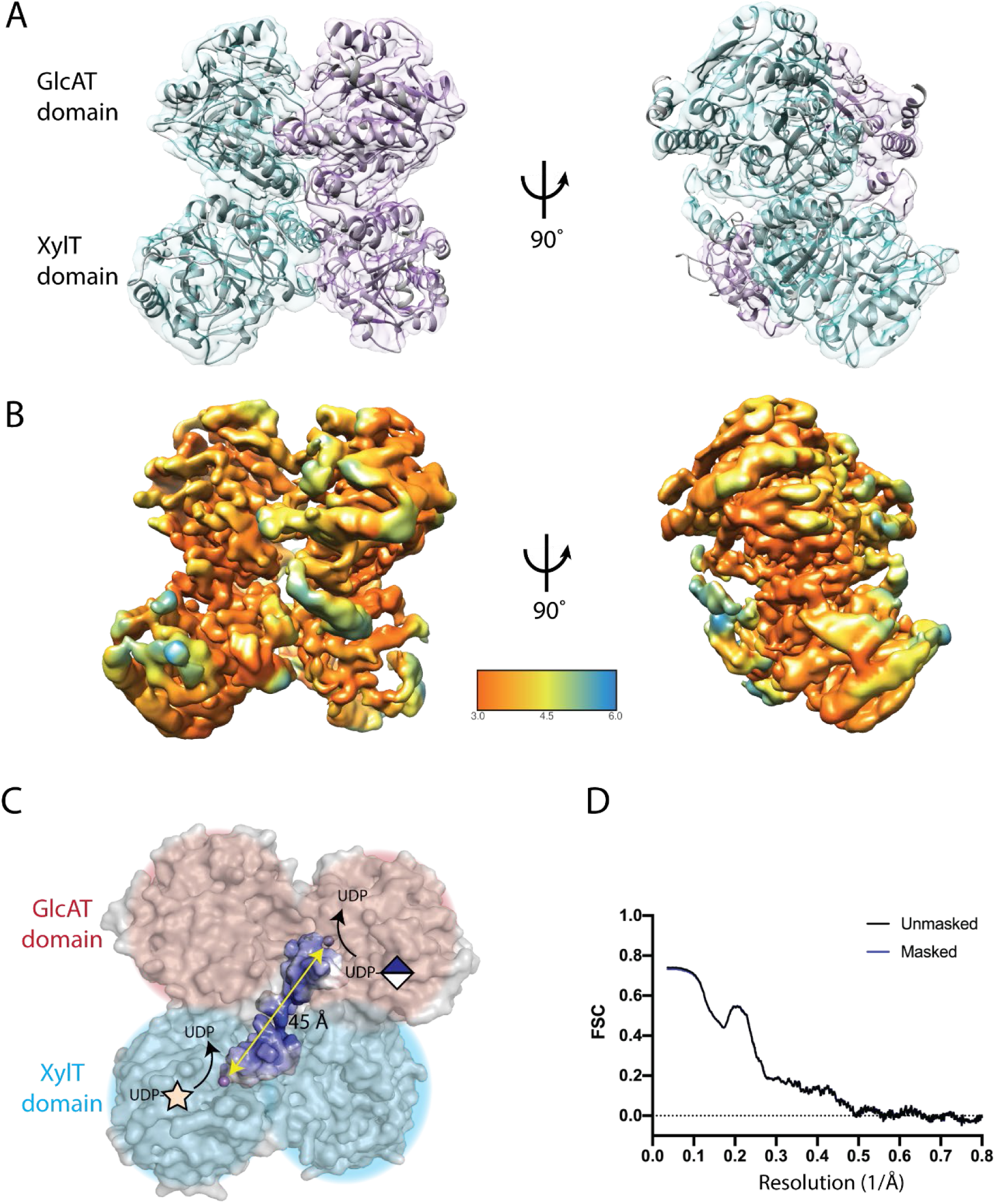
Cryo-EM structure of LARGE1ΔTM. (**A**) Two molecules of LARGE1 (grey cartoon) were refined into the 3.7-Å map of LARGE1ΔTM; protomers are displayed as cyan and purple transparent surfaces. Spheres indicate catalytic manganese ions. (**B**) Reconstructed volume colored by local resolution. (**C**) Orthogonal active sites each coordinating a manganese ion (purple sphere) on alternate protomers face the same direction. A groove (highlighted surface) connects the xylose (cyan shading) and glucuronate (pink shading) transferase active sites on alternate protomers. The quaternary structure implies that matriglycan is polymerized using orthogonal active sites on alternate protomers of LARGE1 (diagonal yellow arrow). (**D**) Map-to-model FSC of LARGE1 catalytic domains’ fit into the reconstructed volume.

The xylose transferase active site is typical in that the manganese ion is chelated by the DXD motif between the β4 and β5 strands and a C-terminal histidine located between β9 and α12. Histidine sidechains appear to coordinate the manganese ion in the glucuronate transferase domain (Fig. S7). The DXD motif in the glucuronate transferase domain are still expected to coordinate the manganese ion during catalysis because the D563N and D565N mutations ablate glucuronate transferase activity (Fig. 3A). The glucuronate transferase domain may have greater plasticity given the structural adjustments required for metal coordination and sugar transfer. This implies that the LARGE1ΔTM cryo-EM volume represents a resting, inactive conformation possibly due to the absence of substrates. LARGE1ΔTM coordinates and maps have been deposited with the following PDB and EMDB accession codes: 7UI6, EMD-26540 (C1 symmetry) and 7UI7, EMD-26541 (C2 symmetry).

**Fig. 3.**
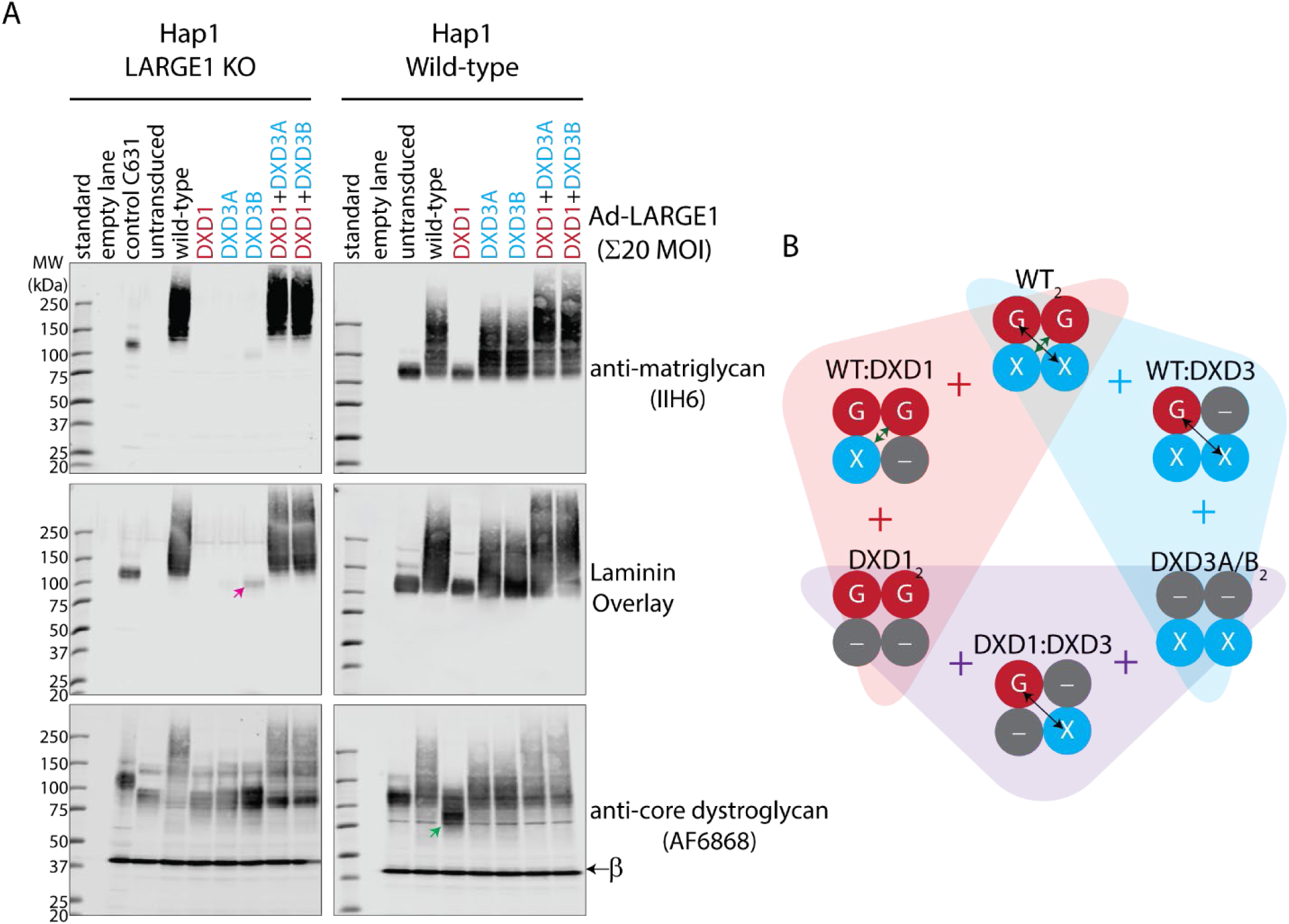
A combination of LARGE1 active site mutants restore matriglycan on native, endogenous dystroglycan. (**A)** Western blot of *LARGE1* KO and wild-type Hap1 cell lines transduced with adenovirus encoding LARGE1 (20 MOI): wild-type (WT), xylose transfer mutant (DXD1; D242N/D244N), glucuronate transfer mutant (DXD3A/B; D563N or D565N, respectively) or a combination of both (DXD1+DXD3A/B). Matriglycan is detected using anti-matriglycan antibody (IIH6) or laminin overlay, whereas core dystroglycan protein is detected using AF6868. Pink arrows indicate a faint matriglycan-positive band in cells transduced with DXD3B in *LARGE1* KO Hap1 cells. Green arrows indicate a perturbation in the migration of dystroglycan in wild-type Hap1 cells transduced with DXD1. (**B**) Adenovirus produce combinations of LARGE1 dimers in Hap1 wild-type or co-transduced *LARGE1* KO cells. Background shading and color-coded plus symbols indicate dimeric combinations present in DXD1 and DXD3 co-transduced *LARGE1* KO Hap1 cells (purple), DXD1 (red) or DXD3 (blue) transduced Hap1 wild-type cells. Double-headed arrows indicate potential processive polymerization of matriglycan by orthogonal active sites on alternate protomers.

The dimeric structure of LARGE1ΔTM reveals that it has two sets of active sites per molecule – a xylose transferase and glucuronate transferase site on each face. Matriglycan is likely polymerized by switching between the active sites on one face (Fig. 2C) rather than between opposite faces, which is less efficient because it would require the non-reducing end of the polymer to navigate around the catalytic head between additions. We suspect the unused transferase domains provide structural support during matriglycan polymerization.

### Structural basis of matriglycan polymerization

The active sites on each protomer in the LARGE1 dimer face opposite directions (Fig. 2) suggesting matriglycan is likely polymerized using the xylosyltransferase site on one protomer with the glucuronate transferase site on the other – that is orthogonal active sites on alternate protomers, which may provide a reason for dimerization. Each glycosyltransferase domain is capable of independently transferring sugars (*1*). We assessed the ability of LARGE1 constructs that were exclusively devoid of either xylose (D242N/D244N; DXD1) or glucuronate (D563N; DXD3A or D565N; DXD3B) transferase activity to synthesize matriglycan on native, endogenous dystroglycan in wild-type and *LARGE1* KO HAP1 cells.

*LARGE1* KO HAP1 cells transduced with adenovirus encoding wild-type LARGE1 restored matriglycan on native, endogenous dystroglycan, whereas LARGE1 mutants lacking either the xylose (DXD1) or glucuronate (DXD3A/B) transferase activity were not able to polymerize matriglycan when transduced in isolation (Fig. 3A). However, a combination of the DXD1 and DXD3A/B LARGE1 mutants synthesized matriglycan on dystroglycan to the same extent as cells transduced with wild-type LARGE1, suggesting that either a DXD1-DXD3 heterodimer or a combination of homodimers (DXD1_2_ and DXD3_2_) can polymerize matriglycan (Fig 3A). The DXD1-DXD3 heterodimer might be capable of polymerizing matriglycan processively because both functions are present in a single entity, whereas a combination of LARGE1 mutant homodimers are obligated to act distributively (Fig. 3B).

Transduction of *LARGE1* KO Hap1 cells with the glucuronate transferase mutant DXD3B (D565N), which retains xylose transfer activity, can produce a faint matriglycan-positive band on Western blots (Fig. 3A; pink arrow). A similar band for the DXD3A (D563N) mutant may be observed when the image contrast is increased (Fig. S9). Therefore, we suspect that the addition of single xylose unit, results in the faint, lower molecular weight band (Fig. 3A; pink arrow). The efficiency of xylose transfer is greater for DXD3B than DXD3A, which suggests that the mutation of aspartate-563 in the glucuronate transferase domain is communicated to the xylose transferase domain better than the mutation of aspartate-565. Thus, we may be observing interdomain communication between the catalytic domains of LARGE1.

Complementing *LARGE1* KO Hap1 cells with wild-type LARGE1 produced matriglycan that appeared dispersed and uncontrolled in length and did not recapitulate the discrete band observed in untransduced control wild-type Hap1 cells (Fig. 3A). This may be due to overexpression of the LARGE1 protein from the adenovirus. Similarly, complementing wild-type Hap1 cells with either of the glucuronate transferase mutants, DXD3A or DXD3B, produced indiscrete matriglycan-positive bands. Conversely, transduction of wild-type Hap1 cells with adenovirus encoding a LARGE1 xylose transferase mutant (DXD1) decreased the size of core dystroglycan (Fig. 3A; green arrow) compared to untransduced controls. A heterodimer, which has at least one DXD1 mutant protomer, might compete for dystroglycan or interfere with matriglycan polymerization by preventing the first xylose transfer.

Heterodimers consisting of a wild-type and a mutant protomer are known to abrogate protein function in a variety of contexts including signaling of G-protein coupled receptors (*21, 22*), DNA binding of basic helix-loop-helix transcriptional factors (*23*), and activation by co-regulators, such as p53 (*24*). Given that isolated homodimers of LARGE1 mutants (DXD1_2_ and DXD3_2_) are incapable of polymerizing matriglycan and that native endogenous LARGE1 is not easily detected by Western blot, we attribute the effects of uncontrolled matriglycan polymerization to the dominant-negative phenotype generated by wild-type mutant heterodimers (wt-DXD1 or wt-DXD3), which would suggest a degree of allostery given that at least one set of orthogonal domains are active.

### Matriglycan of discrete length is processively polymerized on dystroglycan

Biopolymers such as polynucleotides and polypeptides are synthesized using templates that regulate their length and permutation. However, the polymerization of glycans is a template-free process. Branched glycans are produced by the stochastic interaction of precursors and glycosyltransferase (*25*), whereas the length of linear polysaccharides are potentially governed by factors such as competing reactions (*25*), capping of polymers (*26–28*), substrate availability, allosteric modulators (*29–32*) or molecular timers or rulers (*33–36*) that are intrinsic to the enzyme-substrate complex. In the case of matriglycan, processive polymerization is predicated by the tandem arrangement of catalytic domains in LARGE1. Although most polymerases are not purely processive, the discrete distributions of matriglycan-modified dystroglycan as seen in Western blots from various tissues (*11*) suggests that LARGE polymerases may be processive (*37*). We sought to determine whether LARGE1 polymerizes matriglycan predominantly through a distributive or processive mechanism using an *in vitro* assay because *in vivo* co-transduction assays produce a mixture of dimers and because interference from endogenous factors might confound results.

Dystroglycan ordinarily encounters LARGE1 in the Golgi apparatus (Fig. 4A). We wanted a construct of dystroglycan for *in vitro* LARGE1 matriglycan polymerase assays that recapitulate the Golgi enzyme-substrate complex. Therefore, we designed a secreted, soluble construct, DAG1_28-749_^R311A/R312A^, that retains the dystroglycan globular N-terminal domain (DGN) (*38*) to mimic its immature, but relevant, configuration in the Golgi (Fig. 4B). We observed a discrete matriglycan-positive band (150-200 kDa) of recombinant dystroglycan (Fig. 4C) due to the activity of endogenous LARGE1. DAG1_28-749_^R311A/R312A^ without matriglycan (Fig. 4C; supplementary materials) was used as a substrate for *in vitro* matriglycan polymerization assays (Fig. 5).

**Fig. 4.**
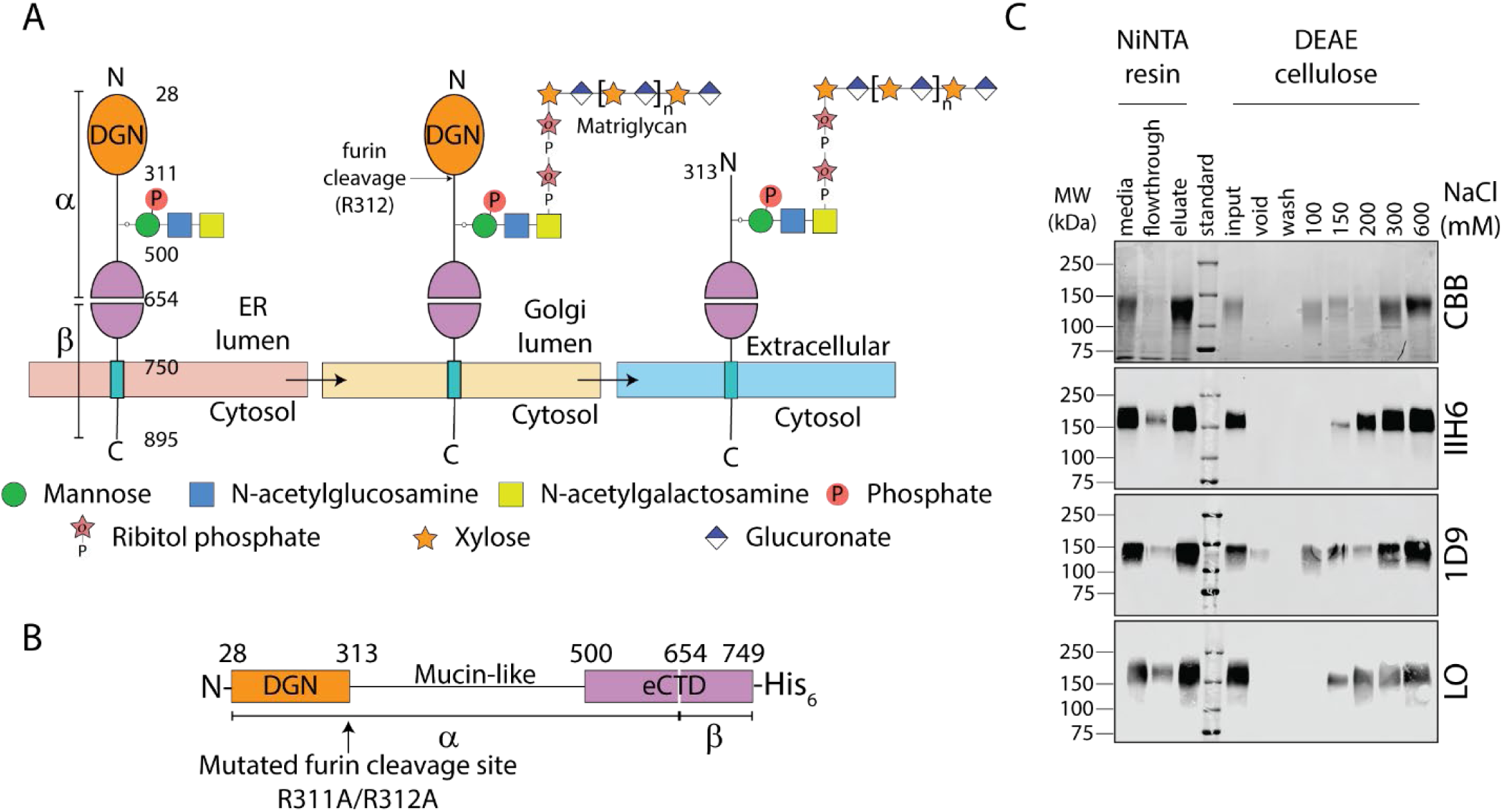
Construct of recombinant Golgi dystroglycan and purification. (**A**) Schematic of dystroglycan glycan modifications through the secretory pathway. The signal peptide (residues 1-27), which is cleaved in the ER, is omitted. Dystroglycan globular N-terminal domain (DGN; orange) and the autoproteolytic extracellular C-terminal domain (eCTD; purple) flank the mucin-like domain (black line). Threonine 317, 319, and 379 in human dystroglycan are post-translationally modified in the ER by POMT1/2, POMGNT2, B3GALNT2, and POMK to make phosphorylated core M3 (N-acetylgalactosamine–β3-N-acetylglucosamine–β4-(phosphate-6-) mannose). In the Golgi, FKTN (FCMD) and FKRP extend core M3 by two phosphoribitol units followed by RXYLT1 (TMEM5) and B4GAT1 which add a xylose-glucuronate primer. Mature dystroglycan lacks DGN. (**B**) Schematic of secreted recombinant Golgi lumenal human dystroglycan retains DGN, which is essential for binding to LARGE1 in the Golgi apparatus, because the furin cleavage site was mutated (R311A/R312A). (**C**) Coomassie brilliant blue (CBB)-stained SDS-PAGE (topmost panel) and Western blot (bottom panels) of recombinant dystroglycan purified from culture media of stably transfected HEK 293 F cells. Protein was eluted using imidazole from IMAC resin and then subject to anion exchange chromatography (DEAE cellulose) to separate dystroglycan carrying matriglycan from unmodified protein. The fraction lacking matriglycan (IIH6) and laminin-binding capacity (LO) were used for matriglycan polymerization assays. The presence of DGN on recombinant dystroglycan was confirmed by Western blotting with 1D9.

**Fig. 5.**
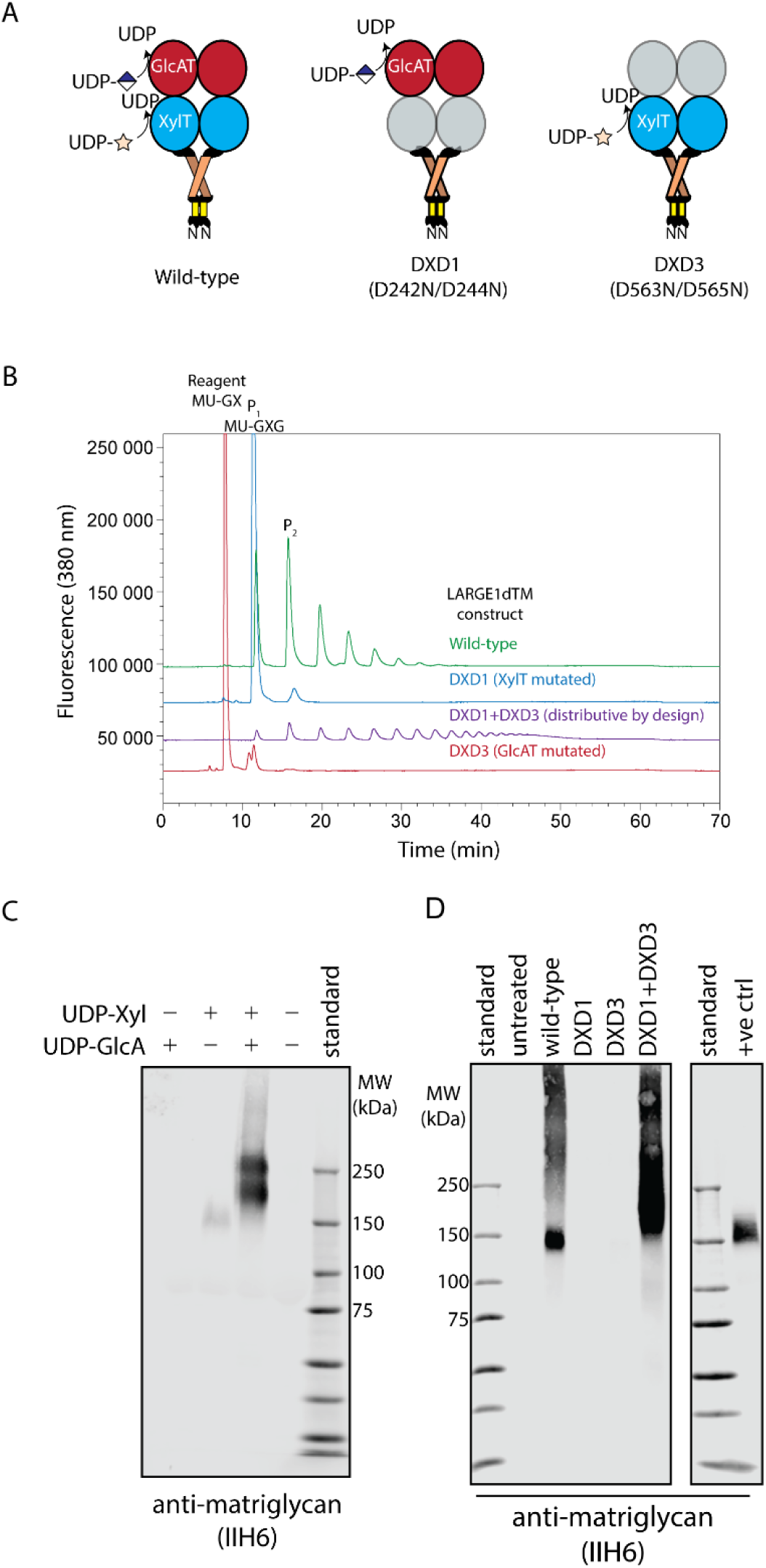
LARGE1 processively polymerizes matriglycan of discrete length on dystroglycan *in vitro*. (**A)** Schematic of LARGE1ΔTM constructs used for determining the mechanism of matriglycan polymerization. Wild-type LARGE1ΔTM construct has both xylose and glucuronate transfer activities whereas DXD1 and DXD3 retain either glucuronate or xylose transfer activities, respectively. (**B**). Anion exchange chromatogram of matriglycan polymerized on 4-methylumbelliferone-glucuronate-xpylose by LARGE1ΔTM constructs. (**C**) Western blot of matriglycan polymerized on recombinant dystroglycan by LARGE1ΔTM in the presence of UDP-sugars at pH 7.4 (n=3). (**D**) Western blot of matriglycan polymerized by LARGE1ΔTM constructs at pH 6.6 (n=3).

Processively synthesized polymers accumulate minimal reaction intermediates (*37*), whereas products synthesized distributively have a Poisson distribution (Fig. 5B and S6). We compared the distribution of matriglycan products on DAG1_28-749_^R311A/R312A^ polymerized by LARGE1ΔTM made by a mixture of the LARGE1ΔTM mutants, DXD1_2_ and DXD3_2_, which are obligated to polymerize matriglycan distributively (Fig. 3B and 5). A discrete matriglycan-positive band on Western blot is absent when a combination of LARGE1ΔTM with either the xylose (DXD_2_) or glucuronate (DXD3_2_) transferase sites mutated polymerize matriglycan, rather we observe a smear of indiscrete lengths (Fig. 5D), which relates to Poisson distribution of products synthesized by LARGE1ΔTM on a synthetic substrate (Fig. 5B and S6). In contrast, polymers synthesized on recombinant dystroglycan by wild-type LARGE1ΔTM produced a discrete band (125-250 kDa), resembling the naturally observed distribution, on a background smear of apparently indiscriminate lengths (Fig. 5C and D). These *in vitro* experiments suggest that a molecular ruler intrinsic to the LARGE1-dystroglycan enzyme-substrate complex controls polymer length, and that LARGE1 primarily polymerizes matriglycan processively.

Interestingly, the faint band observed on Western blots of *LARGE1* KO HAP1 cells transduced with DXD3A/B (Fig. 3A) is also observed in the *in vitro* matriglycan polymerization assay where only UDP-xylose is added as a substrate (Fig. 5C; n = 3). A similarly faint band is observed when protein *O*-mannose kinase (POMK) is inactivated (*29*).

Because standard, ubiquitous GT-A, Rossman-like domains, each with its own catalytic activity, are uniquely arranged in tandem to polymerize a linear chain of alternating sugars, the protein scaffold of LARGE might be amenable to engineering. Potentially, many polymers of alternating monosaccharides could be synthesized by substituting the individual GT-A domains of LARGE. Their lengths may also be controlled by exploiting the molecular “ruler” created by the LARGE1-dystroglycan enzyme-substrate complex. Engineering linear polymers of alternating monosaccharides with defined lengths may be broadly applied in the scientific, medical, or industrial fields.

## Conclusion

The bifunctional glycosyltransferase LARGE is a parallel homodimer. LARGE1 polymerizes matriglycan on dystroglycan using orthogonal active sites on alternate protomers. Matriglycan of discrete length is polymerized processively using a molecular ruler that is likely intrinsic to the LARGE1-dystroglycan enzyme-substrate complex. The standard architecture of tandem Rossmann-like GT-A folds (Fig. S4A) in LARGE proteins may lend themselves to enzyme engineering for the synthesis of polymers of alternating sugars of defined length. The structure and mechanism of LARGE elucidated in this study is an essential first step for the modulation of matriglycan polymerization in cases of neuromuscular disorders as well as LASV and other pathogenic mammarenaviruses that bind matriglycan for cell entry.

## Acknowledgments

We thank Drs. Sandipan Chowdhury, Erhard Hohenester, Andrew Ward, Lance Wells, and Liping Yu for helpful discussions. This research used resources of the Advanced Photon Source, a U.S. Department of Energy (DOE) Office of Science User Facility operated for the DOE Office of Science by Argonne National Laboratory under Contract No. DE-AC02-06CH11357. This project was supported by grants P41 GM103622 and P30 GM138395 from the National Institute of General Medical Sciences of the National Institutes of Health. Use of the Pilatus 3 1M detector was provided by grant 1S10OD018090 from NIGMS. The content is solely the responsibility of the authors and does not necessarily reflect the official views of the National Institute of General Medical Sciences or the National Institute of Health. Extraordinary facility operations were supported in part by the DOE Office of Science through the National Virtual Biotechnology Laboratory, a consortium of DOE national laboratories focused on the response to COVID-19, with funding provided by the Coronavirus CARES Act. A portion of this research was supported by NIH grant U24GM129547 and performed at the PNCC at OHSU and accessed through EMSL (grid.436923.9), a DOE Office of Science User Facility sponsored by the Office of Biological and Environmental Research. We are grateful to Dr. Jennifer Barr of the Scientific Editing and Research Communication Core at the University of Iowa Carver College of Medicine for critical reading of the manuscript. We are grateful to Amber Mower and Jaeda Harmon for assistance with administrative support.

## Author contributions

Conceptualization: SJ, KPC

Methodology: SJ, KPC, NJS, ZX, TY, JH, SC, OD, MEA, DV

Investigation: SJ, KPC, NJS, ZX, JH, TY, JH, SC, OD, MEA, DV

Visualization: SJ, NJS, ZX, TY, MEA, DV

Supervision: KPC

Writing—original draft: SJ, KPC

Writing—review & editing: SJ, KPC, NJS, ZX, TY, MEA, DV

## Data and materials availability

All data are available in the main text or the supplementary materials. LARGE1 coordinates and maps have been deposited with the accession codes 7UI6 and EMD-26540 (C1 symmetry) and 7UI7 and EMD-26541 (C2 symmetry).

## Supplementary Materials

Materials and Methods

Figures S1-S13

Tables S1-S3

