## Supplementary Figures for "Structure and mechanism of LARGE1 matriglycan polymerase"

**This PDF file includes:**

Methods and Materials

Figures S1-S13

Data Tables S1-S3

### Materials and Methods

*Recombinant protein production:* Pure recombinant LARGE proteins were obtained as previously described (*1*). Briefly, LARGE1ΔTM and LARGE2ΔTM secreted by stably transfected HEK 293 F cells cultured in SFM II media supplemented with glutamine and required antibiotics were purified using TALON resin. For recombinant soluble constructs of dystroglycan, HEK 293 F cells were stably transfected (XtremeGENE$^{TM}$ 9 DNA transfection reagent; Sigma-Aldrich) according to the manufacturer's protocol with pcDNA3.1 + encoding human dystroglycan residues 1–749 encoding a mutation in the furin cleavage site (R311A/R312A) and a C-terminal hexahistidine tag. The secreted dystroglycan protein $DAG1_{28–749}^{R311A/R312A}$ was purified using His60 Ni IMAC resin (Takara) and subjected to anion exchange using DEAE resin to obtain a fraction of dystroglycan unmodified by matriglycan. Proteins were snap-frozen in liquid nitrogen and stored at –80 °C.

*Size-exclusion chromatography with on-line multi-angle light scattering and small-angle X-ray scattering (SEC-MALS-SAXS)* was performed at BioCAT beamline 18-ID at the Advanced Photon Source (Argonne National Laboratory, Lemont, IL). Proteins were centrifuged for five minutes at 13,000 rpm. Samples (4–8 mg/mL, 300-500 µL) were applied to a 24 mL Superdex 200 Increase 10/300 GL column (Cytiva) in 20 mM HEPES pH 7.4 and 150 mM NaCl at a flow rate of 0.6 ml/minute using a 1260 Infinity II HPLC (Agilent Technologies). In-line multiangle light scattering (DAWN Helios II) with built-in dynamic light scattering and differential refractive index measurement (Optilab T-rEX) were used to determine weight-average molecular weight (Wyatt Technologies) and degree of glycosylation (ASTRA 8). The sample passed through the UV detector (1260 Infinity II), MALS and then the differential refractometer (Wyatt), in that order, before SAXS.

The SAXS flow cell consisted of a 1.0 mm ID quartz capillary with ~20 µm walls. A buffer sheath was co-flowed to separate sample from the capillary walls and prevent radiation damage (*2*). Scattering intensity was recorded using an Eiger2 XE 9M detector (Dectris) placed 3.67 m from the sample, which provided a q-range of 0.003–0.42 $Å^{-1}$. Half-a-second exposures were acquired every second.

Data were reduced using BioXTAS RAW 2.1.1 (*3*) implementing ATSAS version 3.0.3. Exposures from protein peaks were buffer subtracted using the average of flanking regions. Peaks within the SEC-SAXS elution profile were deconvolved into their components using evolving factor analysis (EFA) with default settings as implemented in RAW (*4*). The validity of the overall deconvolution was assessed on the component concentration profiles and mean error weighted $\chi^2$ for the whole deconvolution range. Individual components were assessed based on the quality of the scattering profile and SAXS-derived molecular weights of the scattering profile compared to the expected values. The forward scattering intensity, I(0), and the radius of gyration ($R_g$) were calculated from the Guinier fitting. The normalized Kratky plot, the pair-distance distribution plot P(r), and the corrected Porod volume were calculated using GNOM (*5*). Low-resolution *ab initio* bead modeling was carried out for all samples using 15 reconstructions from DAMMIF (*6*) in slow mode, averaged by DAMAVER (*7*), and a final structure was refined in DAMMIN (*8*). AMBIMETER was used to assess the ambiguity of reconstructions (*6*). The calculation of theoretical scattering curves for the model was performed by the program CRYSOL (*9*), which also determines the discrepancy ($\chi^2$ value) between the simulated and experimental scattering curves.

The LARGE1ΔTM, LARGE1ΔTM PNGase F-treated, LARGE2ΔTM and LARGE2ΔTM PNGase F-treated SAXS data have been deposited in SASBDB under the accession codes SASDNF8, SASDNG8, SASDNH8 and SASDNJ8, respectively.

*N-glycomics of recombinant LARGE1ΔTM.* 200 μg of LARGE1ΔTM was digested by trypsin in a 50 mM ammonium bicarbonate buffer (pH = 8.4) overnight at 37 °C. N-glycans were released by PNGase F (New England Biolabs) and enriched by C18 solid-phase extraction. Briefly, a C18 Sep-Pak (Waters) cartridge was conditioned by a successive washing of 5 mL methanol, 5 mL ultrapure water, 5 mL acetonitrile, and 15 mL of ultrapure water. The reaction mixture was loaded onto the preconditioned cartridge, and the flow-through was collected and lyophilized. A 1.5 mL slurry of sodium hydroxide in DMSO and 0.6 mL of iodomethane were added to the lyophilized N-glycans. The mixture was vortexed at room temperature for 45 minutes before being extracted by chloroform. The chloroform solution was washed with ultrapure water three times and dried down under a stream of nitrogen. The permethylated N-glycan was dissolved in 50% methanol and loaded onto another preconditioned C18 Sep-Pak cartridge. The cartridge was washed with 5 mL ultrapure water and 3 mL of 15% acetonitrile before eluting by 3 mL of 50% acetonitrile. The purified glycans were lyophilized and dissolved in 10 μL of methanol. One μL of the solution was mixed with 1 μL of 20 mg/mL 3,4-diaminobenzophenone and analyzed by a Bruker ultrafleXtreme MALDI-TOF/TOF.

*Glycoproteomics* 50 μg of LARGE1ΔTM was reduced by 100 μL of 50 mM of dithiothreitol and carbamidomethylated by 100 μL of 100 mM iodoacetamide before dialyzed against 16 L (4 L each time, replace buffer three times) of 50 mM ammonium bicarbonate with a 10 kDa cut-off dialysis cassette (Sigma-Aldrich). The solution was lyophilized and digested by 1 μg MS grade trypsin (Promega) in a 50 mM ammonium bicarbonate buffer (pH=8.4) overnight at 37 °C. The buffer was lyophilized before the tryptic peptides were dissolved in 100 μL of 0.1% formic acid. 2 μL of the solution was injected into EASY-nLC coupled to an orbitrap Fusion Lumos mass spectrometer. A two-solution (solution A: 0.1% formic acid; solution B: 80% acetonitrile, 0.1% formic acid) nanoLC gradient was used to elute an in-house packed C18 column: 3-7% B in 3 minutes, 7-20% B in 60 minutes, 20-42% B in 10 minutes, 42-60% B in 10 minutes, 80-98% B in 10 minutes. The solution B was kept at 98% until 100 minutes. The mass spectrometer was operated under the positive mode. The spray voltage was 1900 V. The ion transfer line temperature was 275 °C. MS resolution was set to 120 K and the mass range was 350-1500 Da. The maximum injection time was 100 ms. The AGC target was 200000. S-lens RF level was 30. MS/MS resolution was 30 K, and the scan started at 80 Da. The maximum injection time was 90 ms. The AGC target was 80000.

The HCD collision energy was 34%. MS was active from 2 to 95 minutes. Glycopeptide spectra were manually analyzed.

*Cryo-EM sample preparation and data acquisition.* LARGE1ΔTM was vitrified on UltrAuFoil 2/2 200 mesh grids in 20 mM PIPES pH 6.6, 150 mM NaCl, 1 mM $MgCl_2$, 1 mM $MnCl_2$ and 1 mM $CaCl_2$ using a Vitrobot Mark IV (FEI). Grids were glow discharged for 60 s, –15 mA on a PELCO easiGlow (Ted Pella) system. Sample (3 µL) was applied to grids in the Vitrobot chamber (24 ºC and 94% humidity) and blotted for three seconds before plunge-freezing in liquid ethane. Data were collected on a Titan Krios G3 microscope (300 kV) using SerialEM with a K3 direct electron detector (Gatan). A total of 5,085 movies were collected at a pixel size of 0.40075 Å/pixel (super-resolution mode) with a dose of ~50 electrons/$A^2$, exposure time of 1.664 seconds, 50 frames, and a defocus range of -0.8 to -2.0 µm.

*Cryo-EM image processing and 3D reconstruction.* Movies were subject to patch motion correction and patch CTF estimation in CryoSPARC (*10*). Initial picks were performed on a subset of data using the blob picker followed by two-dimensional (2D) classification and *ab initio* reconstruction to generate three-dimensional (3D) templates for template-based picking on the full dataset. Particles (1,742,250) were extracted using a 300-pixel box size (0.8015 Å/pixel) and cleaned with multiple rounds of 2D classification. Multiple classes were used for *ab initio* reconstruction followed by heterogeneous refinement to select 81,263 particles for non-uniform refinement (*11*) in C1 (Fig. S5, EMD-26540). Refinement in C2 symmetry was also performed using 107,895 particles (Fig. S5, EMD-26541). Processing was done initially in CryoSPARC version 3.1 and finished in version 3.3.1. Final maps were post-processed using DeepEMhancer (*12*) and used for model building and validation.

*Model building and refinement.* An initial model for LARGE1 was generated by truncating regions from the LARGE1 model in the AlphaFold Protein Structure Database (*13*). The model was initially docked into the density map using Fit in Map in Chimera (*14*). Manual model building was performed in Coot (*15*) and refinement using real-space refinement in Phenix (*16*). Both models (C1, PDB 7UI6 and C2, PDB 7UI7) were subject to comprehensive validation in Phenix. Figures were generated in Chimera and PyMOL. Software used for data processing, model building, and refinement except for CryoSPARC were curated by SBGrid (*17*).

*Adenoviral transduction.* Hap1 cells (wild-type, LARGE1 KO, POMK KO, DAG1 KO and POMK/DAG1 double-KO) were cultured in IMDM containing 2% FCS supplemented with glutamine and penicillin/streptomycin. Cells were transduced with adenovirus-5 (20 MOI) encoding wild-type or mutant LARGE1 D242N/D244N (DXD1), D563N (DXD3A) or D565N (DXD3B). Cells were subsequently cultured in the same media containing 10% FCS for 48 hours at 37 °C.

*Wheat-germ agglutinin (WGA) affinity chromatography.* Cells were washed in PBS and then PBS-containing protease inhibitors (leupeptin, pepstatin A, aprotinin and PMSF). Cells were scraped and incubated in 50 mM Tris pH 7.6, 150 mM NaCl 1% Triton X-100 with protease inhibitors. The solution was rotated for one hour at 4 °C. The supernatant was applied to WGA resin and rotated for 10 minutes. The resin was washed three times with the same buffer containing 0.1% Triton X-100. The slurry was combined with SDS-loading dye and the sample heated for five minutes at 99 °C.

*Matriglycan polymerization assays.* Pure dystroglycan that was devoid of matriglycan or 4-methylimbelliferone-glucuronate-xylose (MU-GX; 0.4 mM) was combined with UDP-xylose (1 mM) and UDP-glucuronic acid (1 mM) with LARGE1ΔTM constructs (wild-type, DXD1 or DXD3) in 20 mM PIPES pH 6.6, 150 mM NaCl, 2 mM $CaCl_2$, 2 mM $MgCl_2$, 2 mM $MnCl_2$ and incubated for 3-16 hours at 37 °C. The reaction in Figure 5C was carried out in buffer with 20 mM HEPES pH 7.4 and 300 mM imidazole instead of PIPES. Reactions were terminated by heating to 99 °C or the addition of SDS-loading dye for substrates 4-methylumbelliferone or recombinant dystroglycan, respectively.

*Anion exchange chromatography.* Samples with matriglycan polymerized on MU-GX were resolved on an anion exchange column (Phenominex SphereClone ™ 5 µm SAXS 80 Å) connected to a HPLC system (Shimadzu Scientific) in an aqueous solution of ammonium phosphate (20-50 mM) at a pH of 6.0 using a gradient of sodium chloride up to 0.5 M. Products were detected by fluorescence of 4-methylumbelliferone using an excitation wavelength 325 nm and an emission wavelength 380 nm.

*SDS-PAGE.* Proteins were resolved on homemade 3-15% polyacrylamide gradient gels in SDS-glycine buffer for either six hours at 200 V or sixteen hours at 60 V at room temperature. The proteins were transferred to PVDF membrane at 800 mA for five hours at 4 °C.

*Western blotting.* Membranes were blocked in either 2% skim milk in 50 mM Tris pH 7.6 and 75 mM NaCl with 0.1% TWEEN 20 (low-salt TBS-T) or fish gelatin dissolved in low-salt TBS-T and incubated with antibodies against matriglycan (IIH6), DGN (1D9) or dystroglycan (AF6868) for 16 h at 4 °C. For laminin overlays membranes were blocked in 5% skim milk dissolved in 10 mM ethanolamine pH 7.6, 140 mM NaCl, 1 mM $MgCl_2$, 1 mM $CaCl_2$. The membranes were overlaid with mouse laminin (ThermoFisher Scientific) in 3% BSA dissolved in the same buffer and incubated for 16 h. The membranes were washed and incubated with rabbit anti-laminin antibody (SigmaAldrich) for 16 h at 4 °C.. Appropriate infra-red fluorescence conjugated secondary antibodies were used in corresponding blotting buffers. The membranes were visualized on a Li-COR Odyssey CLx.

*Statistical analyses.* Statistical analyses are presented in the supplementary materials for method-specific experiments: MALS, SAXS and cryo-EM reconstruction data presented. SEC-MALS-SAXS experiments have been performed multiple times ($n > 3$) over five years with corresponding results. The LARGE1ΔTM volume from single-particle cryo-EM data were independently reconstructed by S.J. and N.J.S, using a 3D classification after the traditional 2D workflow and traditional 2D workflow only, respectively. The independent reconstructions result in equivalent volumes with marginally better resolution for 3D classification only. Matriglycan polymerization probed by Western Blots were performed in triplicate unless otherwise noted.

Transmembrane   Coiled-coil   Mn binding DXD   Mn binding C-His

```
LARGE1 MLGICRGRRKFLAASLSLLCIPAITWIYLFSGSFEDGKPVSLSPLESQAHSPRYTASSQR 60
LARGE2 --MLPRGRPRALGAAALLLLLLLL-GFLLFGGDLGCERREP----GGRAGAPGCFPGP-- 51
          : *** : *.*:  ** :  :  : **.*.:   :        .:* :*    .

       ERESLEVRMREVEEENRALRRQLSLAQGRAPSHRRGNHSKTYSMEEGTGDSENLRAGIVA 120
       -------LM------------------PRVPPDGRLRRAAAL--------DGDPGAGPGD 78
               *                   *.* . * .:: :         . :  **

       GNSSECGQQP-VVEKCETIHVAIVCAGYNASRDVVTLVKSVLFHRRNPLHFHLIADSIAE 179
       HNRSDCGPQPPPPPKCELLHVAIVCAGHNSSRDVITLVKSMLFYRKNPLHLHLVTDAVAR 138
        * *:** **    *** :********:*:****:*****:**:*:****:**::*::*.

       QILATLFQTWMVPAVRVDFYNADELKSEVSWIPNKHYSGIYGLMKLVLTKTLPANLERVI 239
       NILETLFHTWMVPAVRVSFYHADQLKPQVSWIPNKHYSGLYGLMKLVLPSALPAELARVI 198
       :** ***:*********.**:**:** :***********:******** .:***:* ***

       VLDTDITFATDIAELWAVFHKFKGQQVLGLVENQSDWYLGNLWKNHRPWPALGRGYNTGV 299
       VLDTDVTFASDISELWALFAHFSDTQAIGLVENQSDWYLGNLWKNHRPWPALGRGFNTGV 258
       *****:***:**:****:* :*.. *.:***************************:****

       ILLLLDKLRKMKWEQMWRLTAERELMGMLSTSLADQDIFNAVIKQNPFLVYQLPCFWNVQ 359
       ILLRLDRLRQAGWEQMWRLTARRELLSLPATSLADQDIFNAVIKEHPGLVQRLPCVWNVQ 318
       *** **:**:  *********.***:.: :**************::* ** :***.****

       LSDHTRSEQCYRDVSDLKVIHWNSPKKLRVKNKHVEFFRNLYLTFLEYDGNLLRRELFGC 419
       LSDHTLAERCYSEASDLKVIHWNSPKKLRVKNKHVEFFRNFYLTFLEYDGNLLRRELFVC 378
       ***** :*:** :.**************************:***************** *

       PSEADVNSENLQKQLSELDEDDLCYEFRRERFTVHRTHLYFLHYEYEPAADSTDVTLVAQ 479
       PSQPPPGAEQLQQALAQLDEEDPCFEFRQQQLTVHRVHVTFLPHEPP-PPRPHDVTLVAQ 437
       **:   .:*:**: *::***:* *:***::::****.*: ** :*        *******

       LSMDRLQMLEAICKHWEGPISLALYLSDAEAQQFLRYAQGSEVLMSRHNVGYHIVYKEGQ 539
       LSMDRLQMLEALCRHWPGPMSLALYLTDAEAQQFLHFVEASPVLAARQDVAYHVVYREGP 497
       ***********:*:** **:******:********::.:.* ** :*::*.**:**:**

       FYPVNLLRNVAMKHISTPYMFLSDIDFLPMYGLYEYLRKSVIQLDLANTK-KAMIVPAFE 598
       LYPVNQLRNVALAQALTPYVFLSDIDFLPAYSLYDYLRASIEQLGLGSRRKAALVVPAFE 557
       :**** *****: :  ***:********* *.**:*** *: **.*.. :  *::*****

       TLRYRLSFPKSKAELLSMLDMGTLFTFRYHVWTKGHAPTNFAKWRTATTPYRVEWEADFE 658
       TLRYRFSFPHSKVELLALLDAGTLYTFRYHEWPRGHAPTDYARWREAQAPYRVQWAANYE 617
       *****:***:**.***::** ***:***** * :*****::*:** * :****:* *::*

       PYVVVRRDCPEYDRRFVGFGWNKVAHIMELDVQEYEFIVLPNAYMIHMPHAPSFDITKFR 718
       PYVVVPRDCPRYDPRFVGFGWNKVAHIVELDAQEYELLVLPEAFTIHLPHAPSLDISRFR 677
       ***** ****.** *************:***.****::***:*: **:*****:**::**

       SNKQYRICLKTLKEEFQQDMSRRYGFAALKYLTAENNS------ 756
       SSPTYRDCLQALKDEFHQDLSRHHGAAALKYLPALQQPQSPARG 721
       *.  ** **::**:**:**:**::* ****** * ::
```

**Fig. S1. Sequence alignment of human LARGE1 and LARGE2**. Human LARGE1 and 2 were aligned using Clustal Omega to show high sequence identity (*) and similarity (:) of the catalytic domains. The coiled-coil domain (green highlight) is absent in LARGE2.

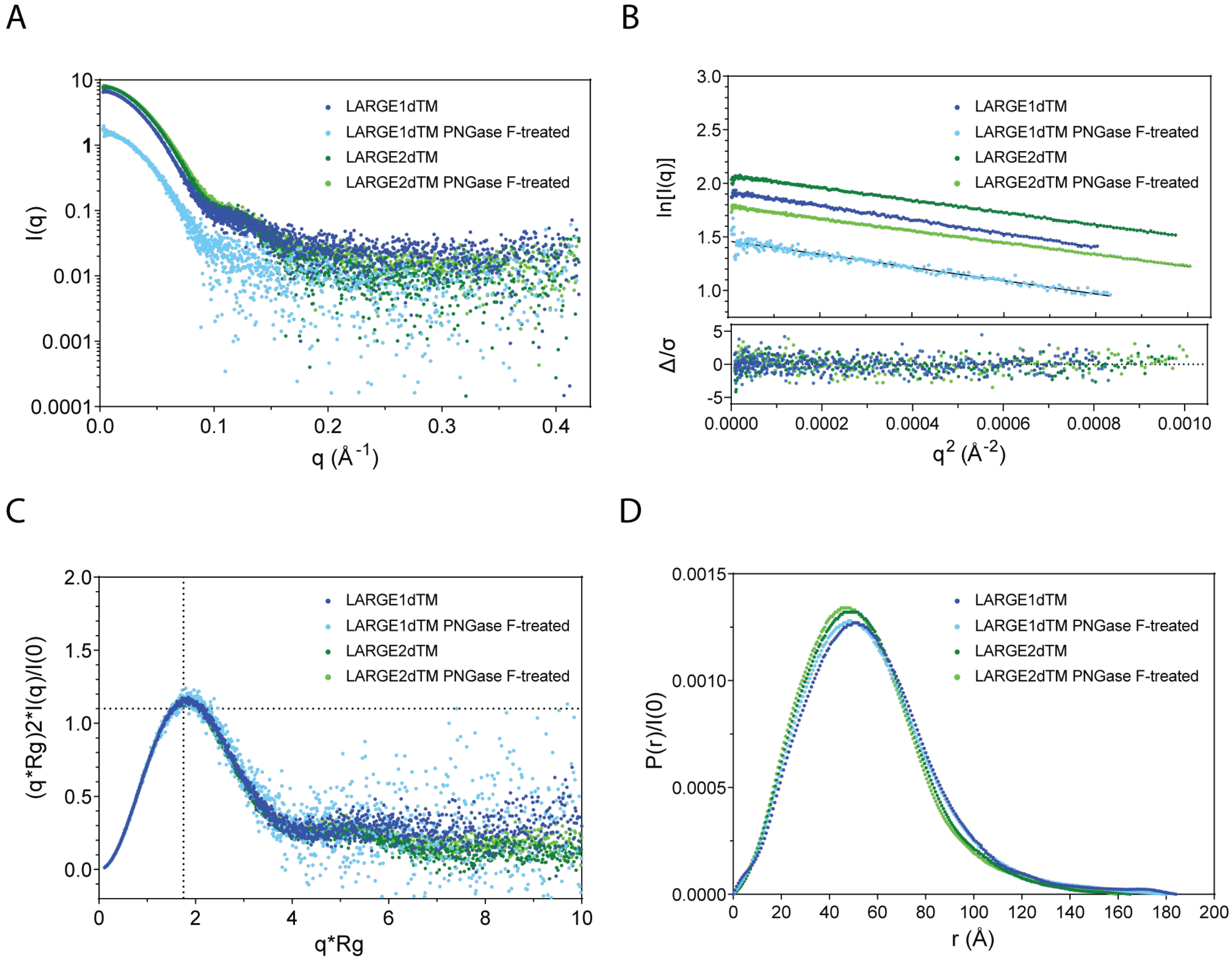


**Fig. S2. Small-angle X-ray Scattering (SAXS) data for LARGE proteins.** (**A**) Scattering intensity (I) over radial distance (q). (**B**) Guinier plots showing linear distribution at low-q (upper panel) and residuals (lower panel), suggesting a lack of interparticle interference. (**C**) Normalized Kratky plot showing globular well-folded entities. (**D**) Histogram of interatomic distance vectors. The maximum dimension is ~160–180 Å.

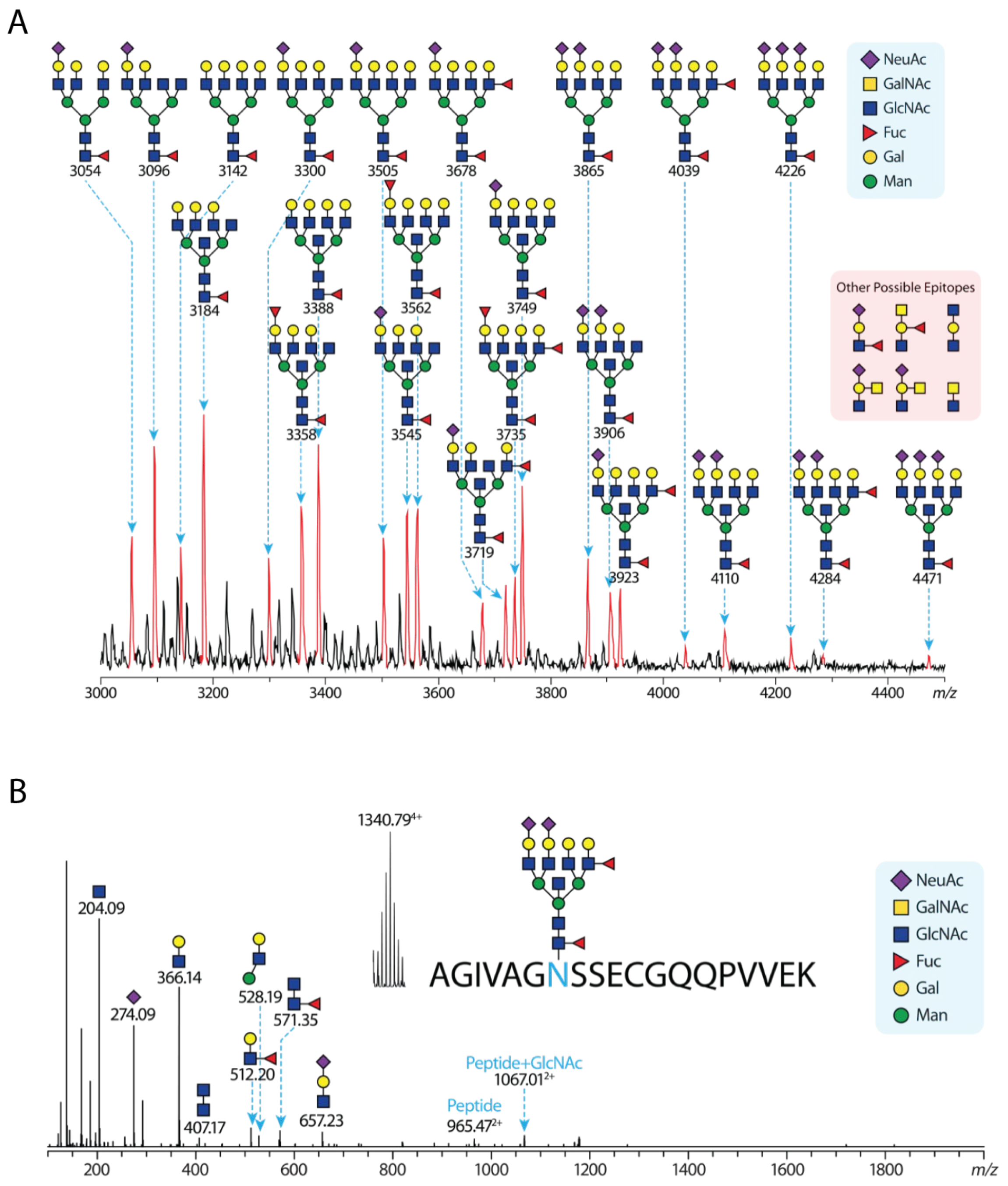


**Fig. S3. Mass spectrometry analysis of LARGE1ΔTM N-glycosylation (N-glycopeptide) (A)** Glycoproteomics of LARGE1ΔTM. (**B**) A representative MSMS spectrum of LARGE1ΔTM N-glycopeptide. Structural annotation of glycan is not unambiguous. MALDI-TOF profiling of complex N-glycans released from LARGE1ΔTM. Sodiated permethylated N-glycans were observed. Structural annotation is not unambiguous and is in favor of more commonly known N-glycan structures.

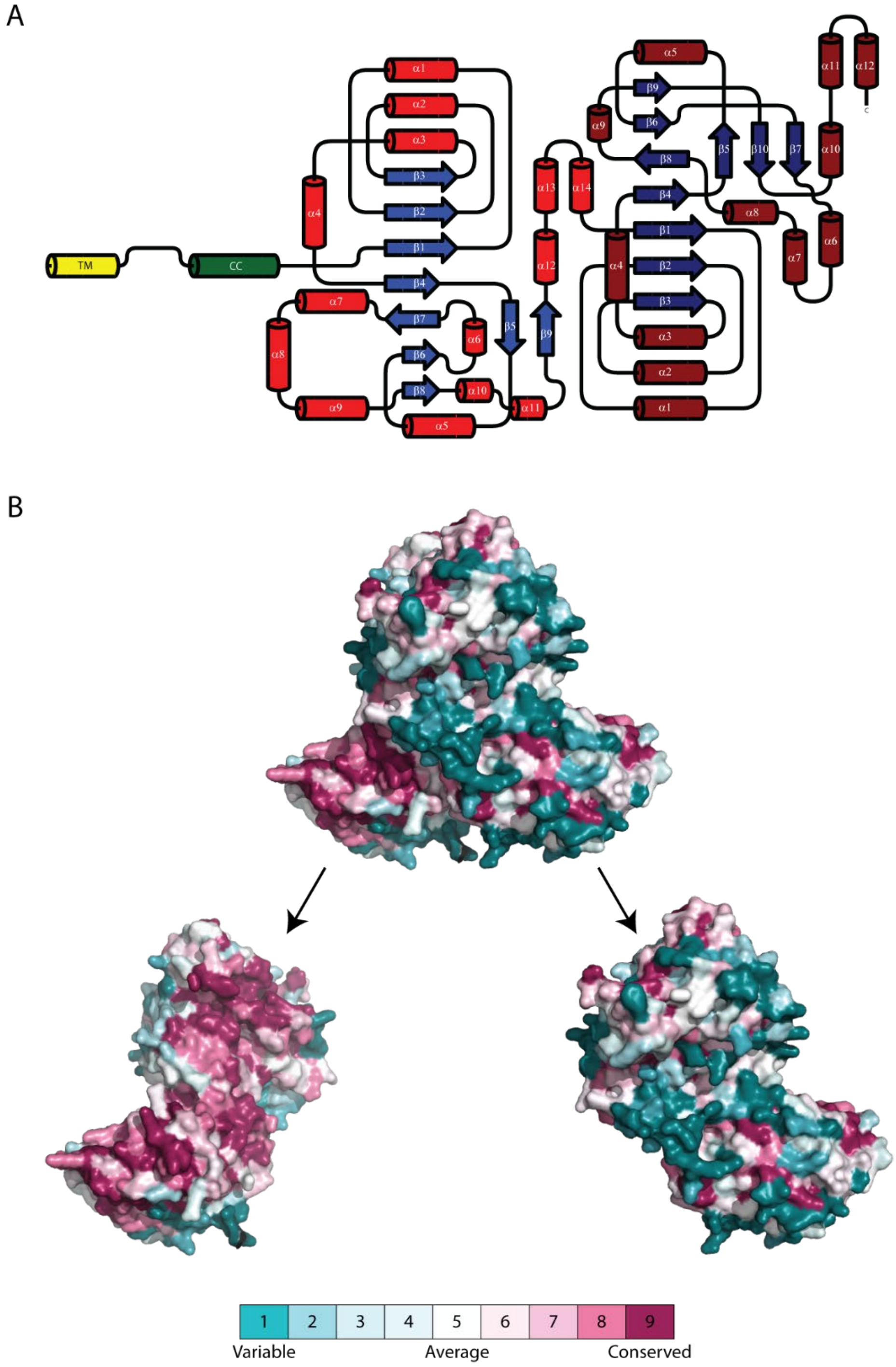


**Fig. S4. Architecture and conservation of LARGE1.** (**A**) The secondary structure of LARGE1 showing α-helices (cylinders) and β-strands (blue arrows), transmembrane (TM; yellow) and coiled-coil (CC; green) and tandem Rossman-folds are depicted. (**B**) The dimerization interface of LARGE1 is conserved (ConSurf; (*18–20*)). Upper panel shows the dimer, whereas the lower panel shows separate protomers in the same orientation, which exposes the highly conserved dimerization interface on the posterior protomer (left).

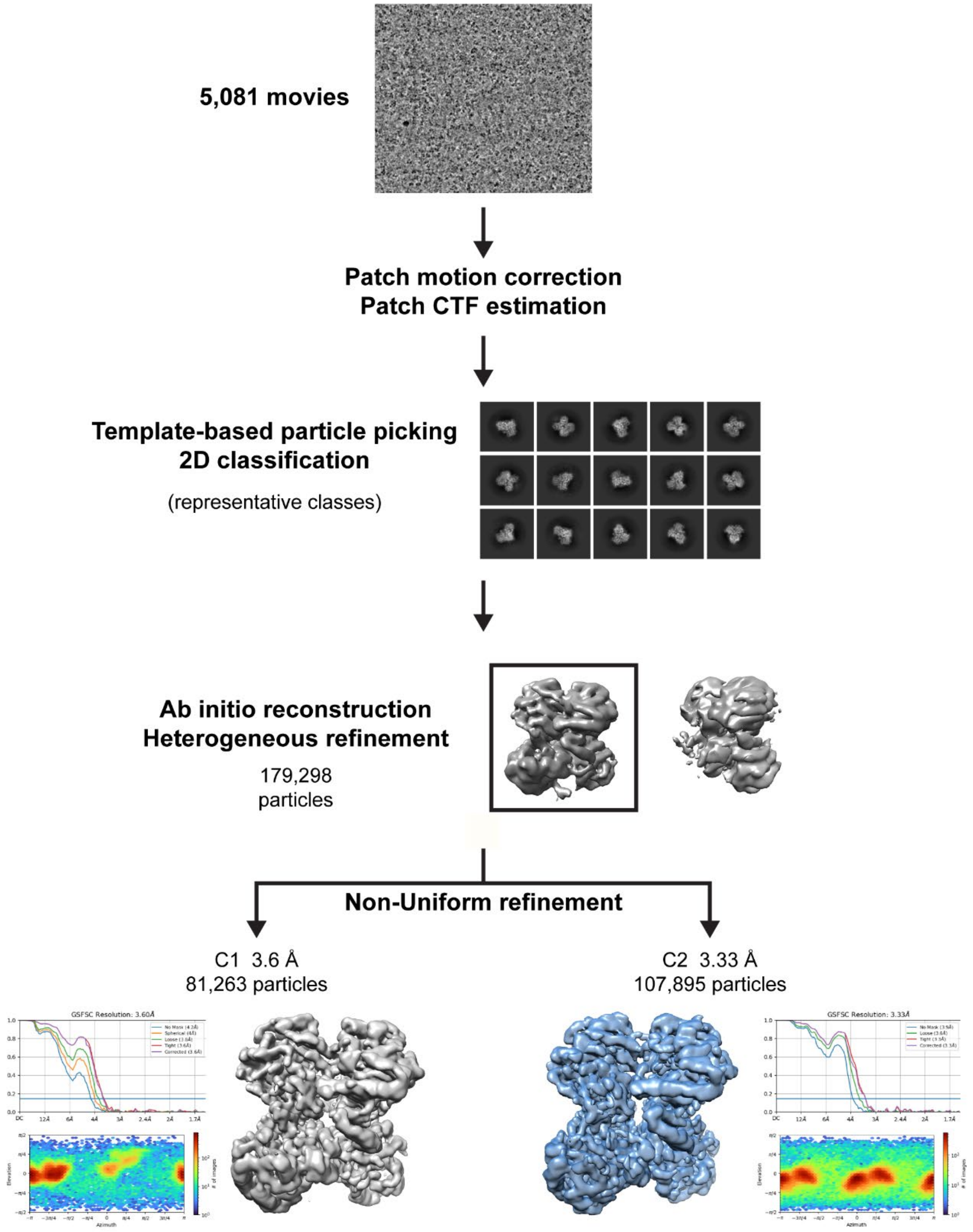


**Fig. S5. LARGE1ΔTM cryo-EM processing workflow.** Cryo-EM reconstruction of LARGE1ΔTM in CryoSPARC. Templates-picker was used on motion- and CTF-corrected micrographs of LARGE1ΔTM. Sample of selected 2D class averages used for *ab initio* reconstruction. Particles were non-uniformly refined separately in C1 and C2 symmetries.

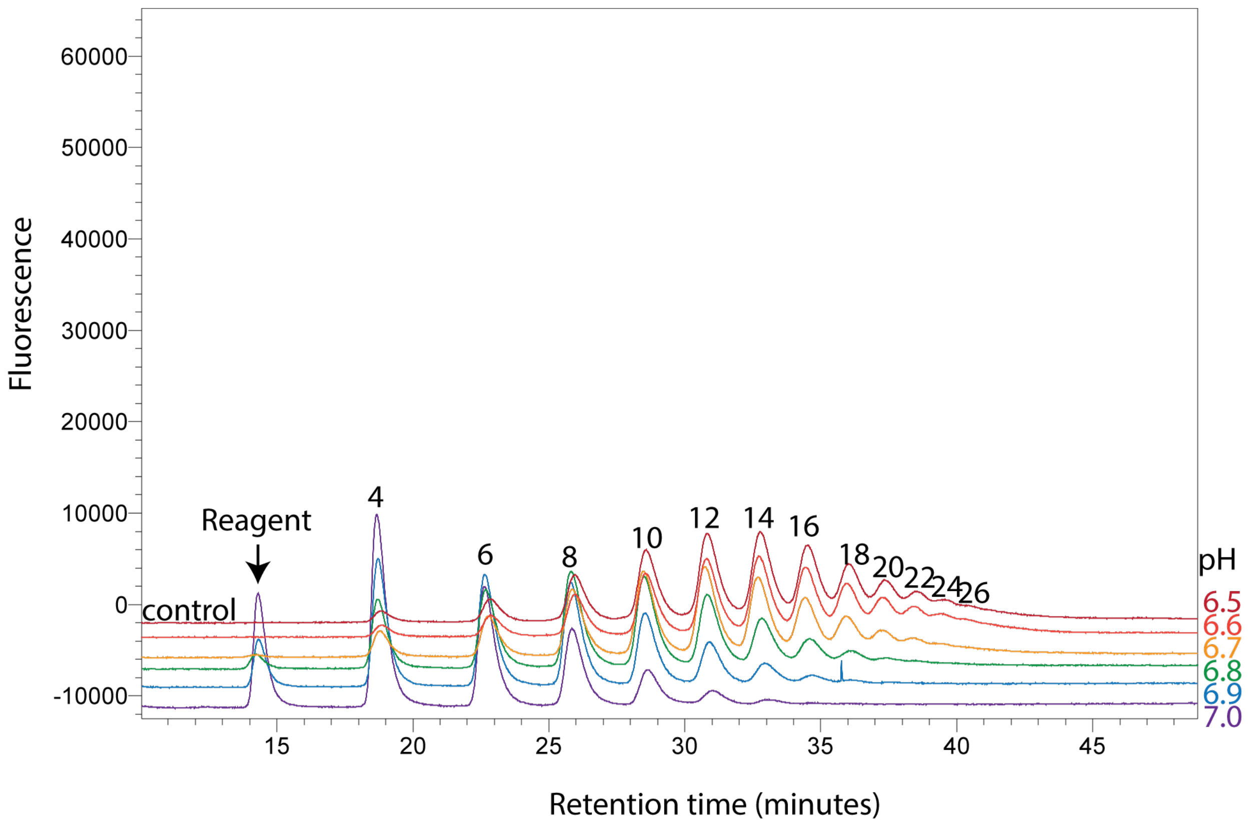


**Fig. S6. Anion exchange chromatogram of matriglycan polymerized by LARGE1ΔM on 4-methylumbelliferyl-glucuronate-xylose.** Product length (enumerated peaks represent the number of monosaccharides) is inversely proportional to the pH of the reaction and product abundance follows a Poisson distribution, which is characteristic of distributive polymerization.

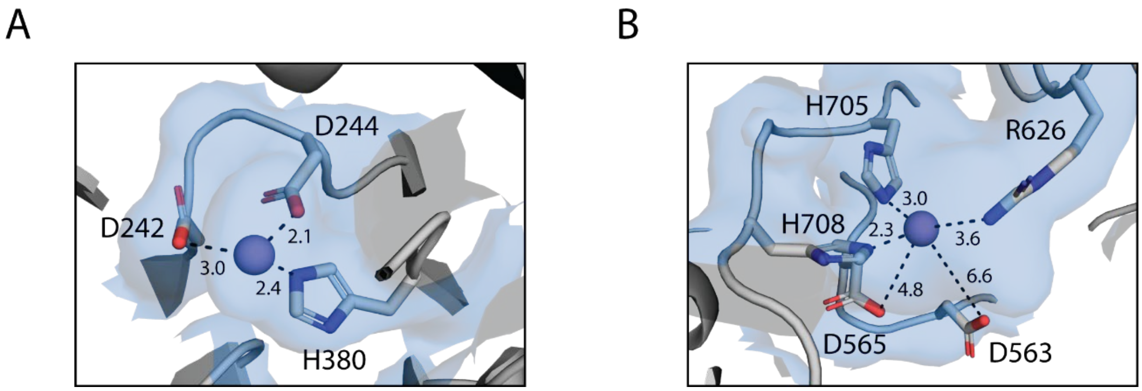


**Fig. S7. LARGE1 active sites.** Manganese ions are coordinated by aspartate residues contributed from DXD motifs and a C-terminal histidine residue in xylose transferase (**A**) but not glucuronate transferase (**B**) active site.

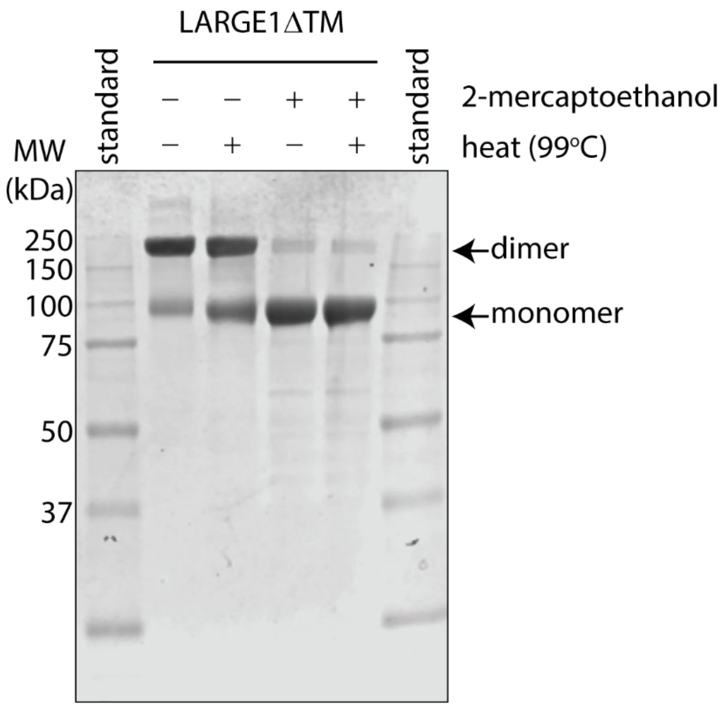


**Fig. S8. Reducing agent monomerizes partially SDS-resistant LARGE1ΔTM dimers.** SDS-PAGE of LARGE1ΔTM treated with SDS with or without reducing agent (2-mercaptoethanol) and heated at 99 °C for five minutes.

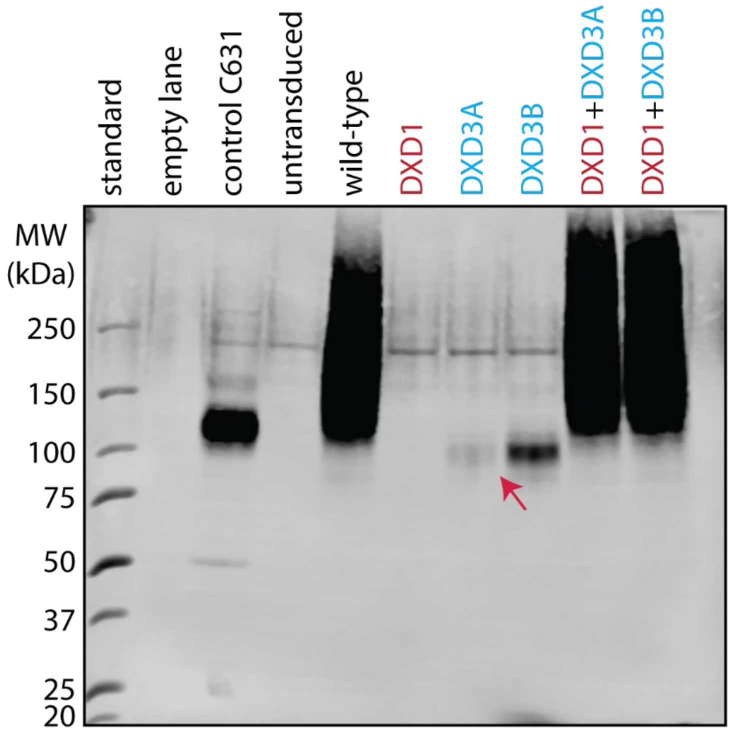


**Fig. S9. Laminin overlay at higher contrast.** Pink arrow shows a very faint band for DXD3A.

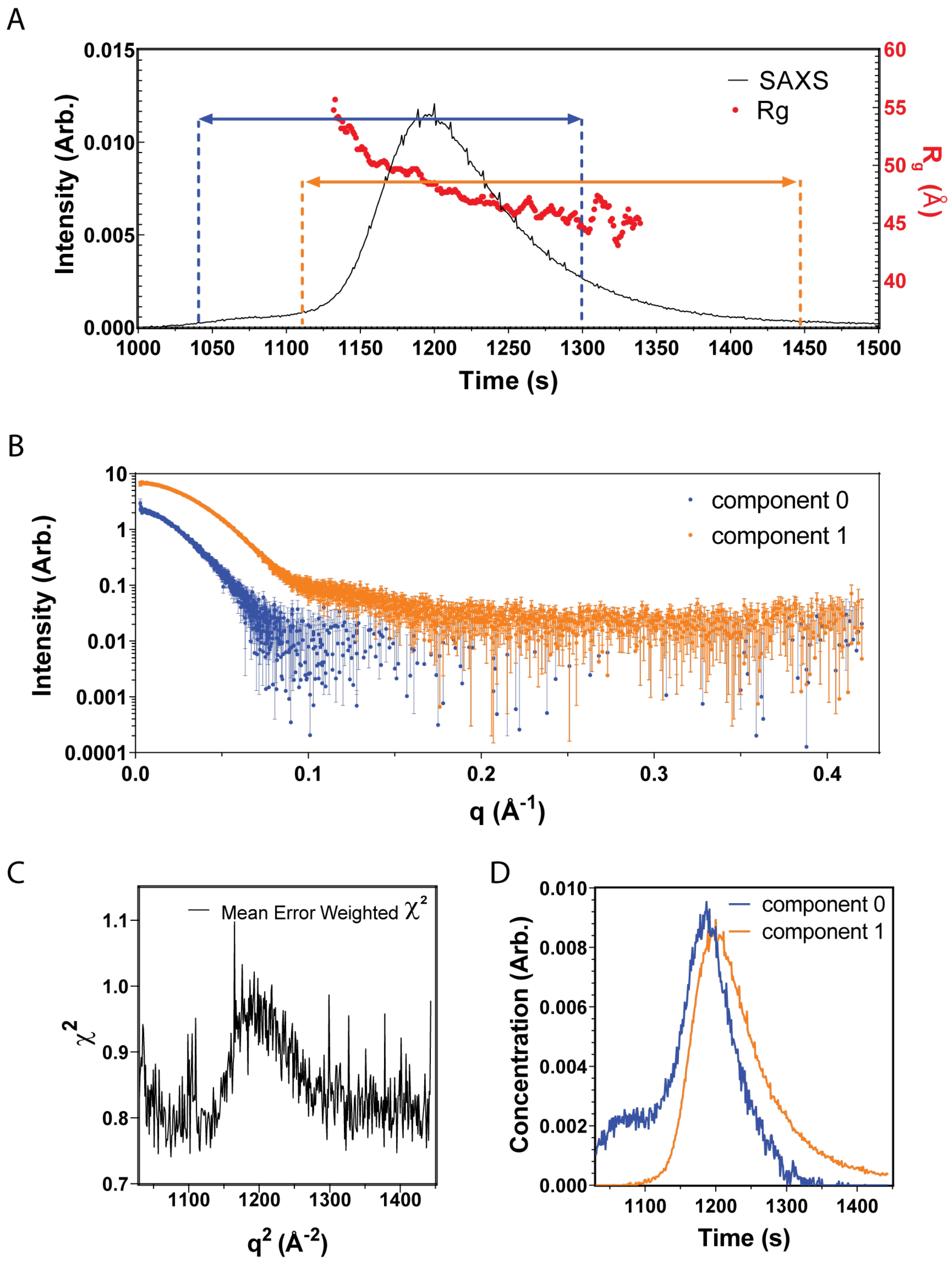


**Fig. S10. SAXS EFA of LARGE1ΔTM.** (**A**) The buffer subtracted integrated SAXS intensity (left axis, in arbitrary scale) and calculated $R_g$ (right axis) as a function of time for the SEC-SAXS experiment. The regions denoted by dashed lines and arrows were deconvoluted by evolving factor analysis (EFA). (**B**) Scattering profiles for components determined by EFA. (**C**) Mean error-weighted $\chi^2$ of the EFA deconvolution. (**D**) Area-normalized component concentration profiles determined by EFA. Colors correspond to component colors on other panels.

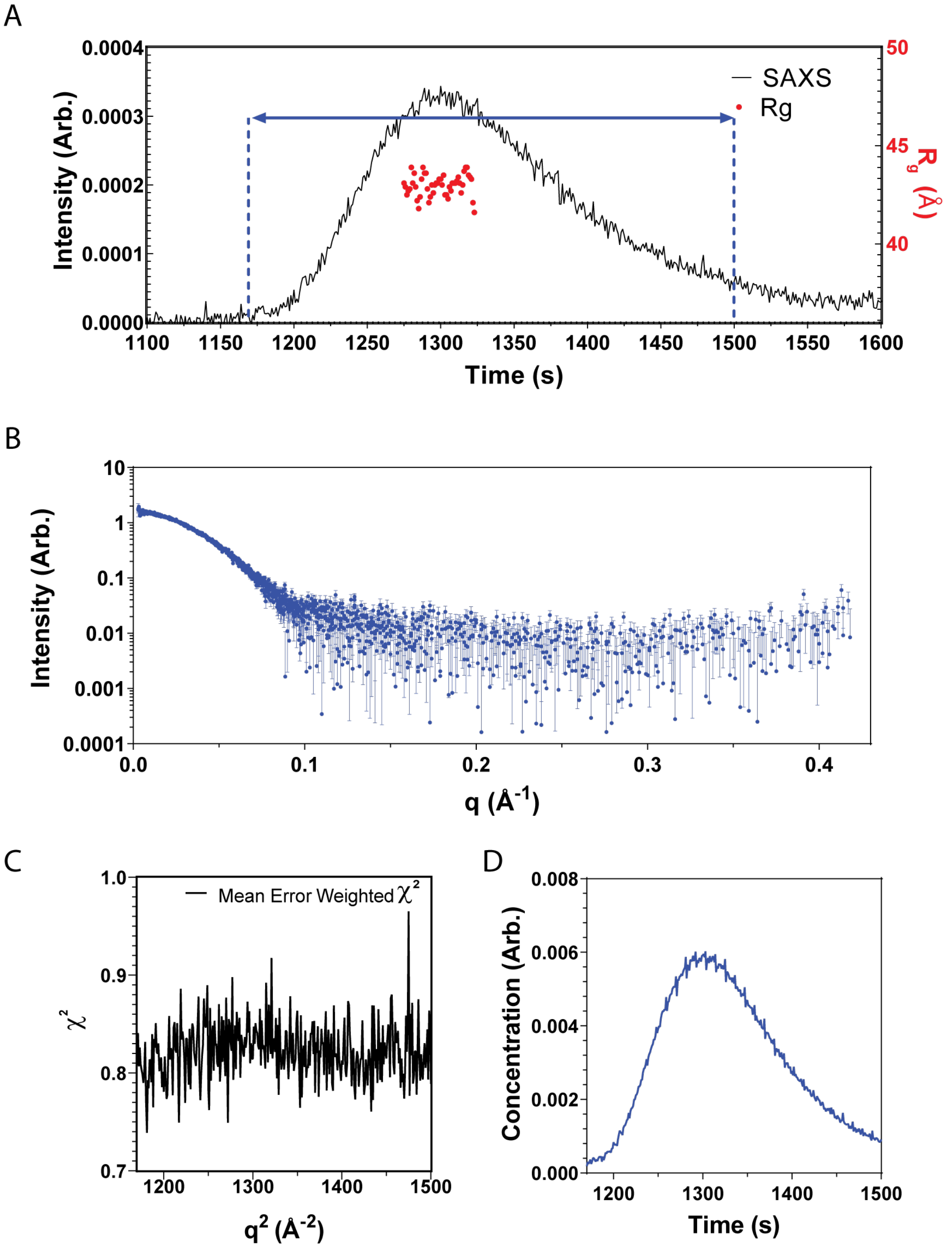


**Fig. S11. SAXS EFA of LARGE1ΔTM treated with PNGase F.** (**A**) The buffer subtracted integrated SAXS intensity (left axis, in arbitrary scale) and calculated $R_g$ (right axis) as a function of time for the SEC-SAXS experiment. The regions denoted by dashed lines and arrows were deconvoluted by evolving factor analysis (EFA). (**B**) Scattering profiles for components determined by EFA. (**C**) Mean error-weighted $\chi^2$ of the EFA deconvolution. (**D**) Area-normalized component concentration profiles determined by EFA. Colors correspond to component colors on other panels.

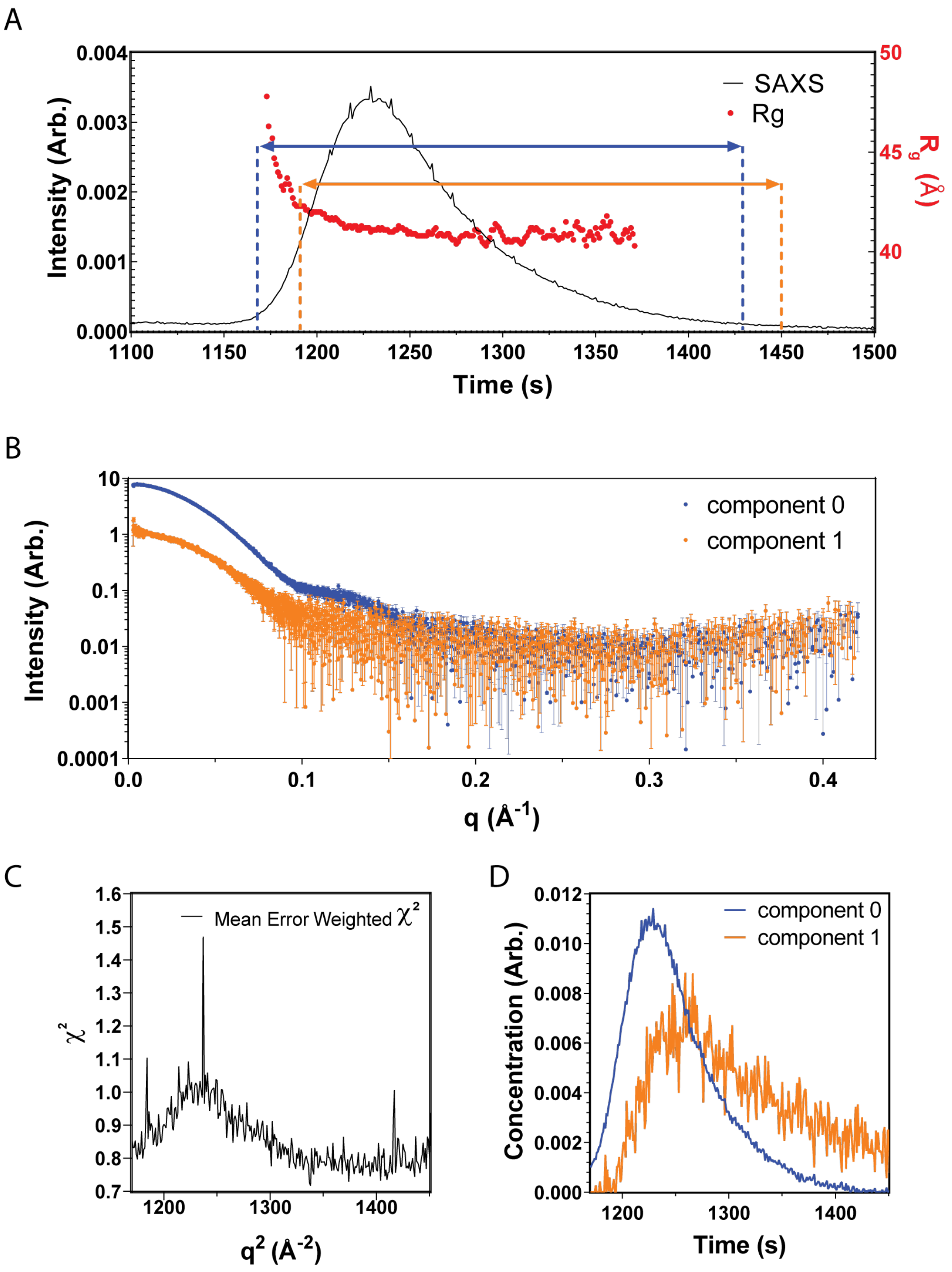


**Fig. S12. SAXS EFA of LARGE2ΔTM.** (**A**) The buffer subtracted integrated SAXS intensity (left axis, in arbitrary scale) and calculated $R_g$ (right axis) as a function of time for the SEC-SAXS experiment. The regions denoted by dashed lines and arrows were deconvoluted by evolving factor analysis (EFA). (**B**) Scattering profiles for components determined by EFA. (**C**) Mean error-weighted $\chi^2$ of the EFA deconvolution. (**D**) Area-normalized component concentration profiles determined by EFA. Colors correspond to component colors on other panels.

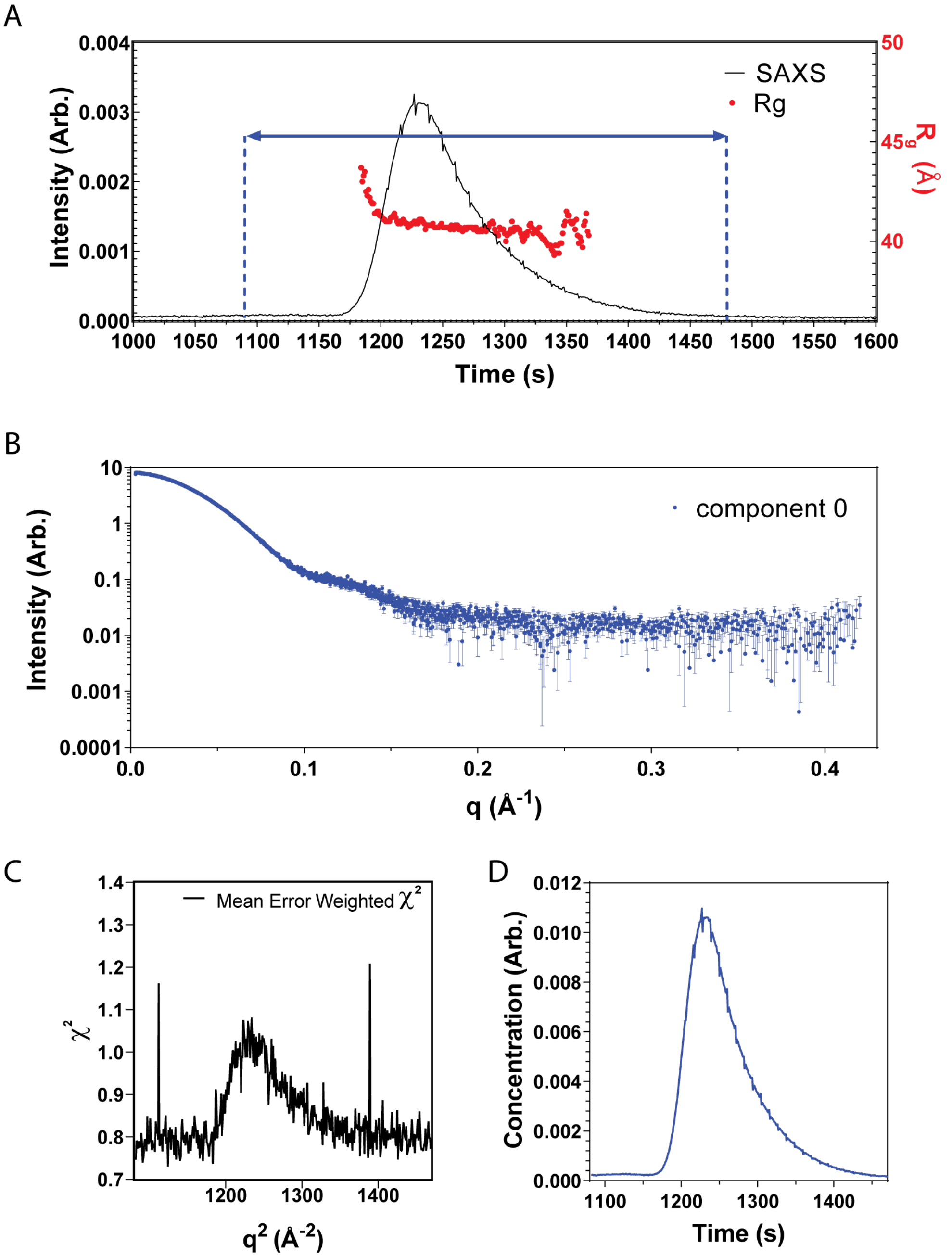


**Fig. S13. SAXS EFA of LARGE2ΔTM treated with PNGase F.** (**A**) The buffer subtracted integrated SAXS intensity (left axis, in arbitrary scale) and calculated $R_g$ (right axis) as a function of time for the SEC-SAXS experiment. The regions denoted by dashed lines and arrows were deconvoluted by evolving factor analysis (EFA). (**B**) Scattering profiles for components determined by EFA. (**C**) Mean error-weighted $\chi^2$ of the EFA deconvolution. (**D**) Area-normalized component concentration profiles determined by EFA. Colors correspond to component colors on other panels.

**Table S1.** SAXS data collection and analysis parameters

| | |
|---|---|
| **a) Sample details** | |
| SEC Column | Superdex 200 increase 10/300 |
| Loaded concentration (mg/ml) | 4-8 |
| Injection volume (ul) | 300-500 |
| Flow rate (ml/min) | 0.6 |
| Solvent (solvent blanks taken from SEC flowthrough prior to elution of protein) | 20 mM HEPES pH 8.0, 150 mM NaCl |
| **b) SAXS data-collection parameters** | |
| Instrument | BioCAT facility at the Advanced Photon Source beamline 18ID with Eiger2 XE 9M (Dectris) detector |
| Wavelength (Å) | 1.033 |
| Beam size (um$^2$) | 150 (h) x 25 (v) focused on the detector |
| Camera length (m) | 3.6 |
| $q$ measurement range (Å$^{-1}$) | 0.003-0.42 |
| Absolute scaling method | Glassy Carbon, NIST SRM 3600 |
| Basis for normalization to constant counts | To incident intensity, by ion chamber counter |
| Monitoring for radiation damage | Automated frame-by-frame comparison of relevant regions using CORMAP (Franke et al., 2015) implemented in BioXTAS RAW |
| Exposure time | 0.5 s exposure time with a 1 s total exposure period (0.5 s on, 0.5 s off) of entire SEC elution |
| Sample configuration | SEC-MALS-SAXS. Size separation used a Superdex 200 Increase 10/300 GL column and a 1260 Infinity II HPLC (Agilent Technologies). UV data were measured using the HPLC, and MALS-DLS-RI data by DAWN HELEOS-II (17 MALS + 1 DLS channels) and Optilab T-rEX (RI) instruments (Wyatt Technology). SAXS data was measured in a sheath-flow cell with an effective path length of 0.542 mm. |
| Sample temperature (ºC) | 23 |
| **c) Software employed for SAXS data reduction, analysis, and interpretation** | |
| SAXS data reduction | Radial averaging; frame comparison, averaging, and subtraction done using BioXTAS RAW 2.1.1 (Hopkins et al., 2017) |
| Basic analysis: Guinier, MW, Normalized Kratky, P(r) | Guinier fit and M.W. using BioXTAS RAW, P(r) function using GNOM (Svergun, 1992). RAW uses MoW and Vc M.W. methods (Rambo & Tainer, 2013; Piiadov et al., 2018) |
| MALS-DLS-RI analysis | Astra 8 (Wyatt) |

**Table S2**. Data collection and model refinement statistics.

| | **LARGE1 C1 symmetry** | **LARGE1 C2 symmetry** |
|---|---|---|
| ***Data collection and image processing*** | | |
| Microscope | Titan Krios | Titan Krios |
| Voltage (kV) | 300 | 300 |
| Camera | K3 | K3 |
| Magnification | 29,000 x | 29,000 x |
| Electron exposure ($e^-/Å^2$) | 50 | 50 |
| Exposure time (s) | 1.66 | 1.66 |
| Number of frames | 50 | 50 |
| Defocus range (µm) | -0.8 to -2.0 | -0.8 to -2.0 |
| Super resolution pixel size (Å) | 0.40075 | 0.40075 |
| Number of movies | 5,085 | 5,085 |
| Particles for final reconstruction (no.) | 81,263 | 107,895 |
| Map resolution (Å)<br>Half map FSC threshold | 3.7<br>0.143 | 3.4<br>0.143 |
| EMDB accession code | 26540 | 26541 |
| ***Model refinement statistics*** | | |
| *Model composition*<br>Non-hydrogen atoms<br>Protein residues<br>Ligands | <br>10,148<br>609<br>4 | <br>10,214<br>613<br>4 |
| *Cross correlation*<br>Mask<br>Volume | <br>0.58<br>0.59 | <br>0.66<br>0.67 |
| *RMSD*<br>Bond length (Å)<br>Bond Angles (°) | <br>0.005<br>0.971 | <br>0.004<br>0.612 |
| *Ramachandran*<br>Favored (%)<br>Allowed (%)<br>Outlier (%) | <br>94.96<br>4.79<br>0.25 | <br>92.36<br>7.64<br>0.00 |
| *Validation*<br>MolProbity score<br>Clashscore<br>Rotamer outliers (%) | <br>2.15<br>18.99<br>1.09 | <br>2.13<br>14.09<br>0.00 |
| PDB accession code | 7UI6 | 7UI7 |

**Table S3**: SAXS analysis structural parameters and *ab initio* model fitting statistics

| **(a) Structural parameters** | | | | |
|---|---|---|---|---|
| | LARGE1dTM | LARGE1dTM PNGase F treated | LARGE2dTM | LARGE2dTM PNGase F treated |
| Guinier Analysis | | | | |
| I(0) (Arb.) | 6.818 ± 0.007 | 1.584 ± 0.004 | 7.951 ± 0.005 | 8.018 ± 0.004 |
| $R_g$ (Å) | 43.99 ± 0.08 | 42.83 ± 0.12 | 41.56 ± 0.05 | 40.87 ± 0.03 |
| q-range (Å$^{-1}$) | 0.00391-0.02946 | 0.00428-0.03033 | 0.00279-0.0312 | 0.00279-0.03169 |
| $q_{max}R_g$ | 1.2959 | 1.2991 | 1.2966 | 1.2952 |
| Coefficient of correlation, $r^2$ | 0.9973 | 0.9752 | 0.9957 | 0.9984 |
| Volume (Å$^3$ adjusted $V_P$ as SAXS MoW2) | 264425 | 248999 | 256585 | 230571 |
| MW, MoW2 method (kDa) | 219 (1.22) | 206 (1.15) | 212 (1.26) | 191 (1.13) |
| MW, Vc method (kDa) | 173 (0.96) | 173 (0.96) | 184 (1.09) | 161 (0.96) |
| *P*(*r*) analysis | | | | |
| I(0) (Arb.) | 6.845 ± 0.010 | 1.595 ± 0.005 | 7.973 ± 0.006 | 8.048 ± 0.004 |
| $R_g$ (Å) | 45.05 ± 0.16 | 44.22 ± 0.29 | 42.18 ± 0.08 | 41.66 ± 0.04 |
| $D_{max}$ (Å) | 184 | 180 | 165 | 159 |
| q-range (Å$^{-1}$) | 0.00391-0.4202 | 0.00428-0.4202 | 0.00279-0.4202 | 0.00279-0.4202 |
| $\chi^2$ (total estimate from GNOM) | 0.907 (0.711) | 2.078 (0.7647) | 1.051 (0.7729) | 1.641 (0.8151) |

| **(b) Shape modelling fitting parameters** | | | | |
|---|---|---|---|---|
| DAMMIF (Slow mode, default parameters, 15 calculations | | | | |
| q-range for fitting (Å$^{-1}$) | 0.00391-0.18185 | 0.00428-0.18669 | 0.00279-0.19258 | 0.00279-0.19587 |
| Symmetry, anisotropy assumptions | P1, unknown | P1, unknown | P1, unknown | P1, unknown |
| Ambiguity score (AMBIMETER) | 0.4771 | 1.505 | 0.4771 | 0.4771 |
| NSD (standard deviation), No. of clusters | 0.805 (0.042), 1 | 0.834 (0.053), 1 | 0.690 (0.074), 1 | 0.703 (0.061), 1 |
| $\chi^2$ range | 1.401-1.476 | 4.258-4.343 | 2.491-2.951 | 6.646-7.315 |
| Model MW estimate range (ratio to expected) | 216.4 (1.21) | 239.8 (1.34) | 242.1 (1.44) | 221.5 (1.32) |
| Model Rg range | 45.161-45.186 | 46.39-46.42 | 42.618-42.644 | 42.186-42.29 |
| Model Dmax range | 180-202 | 179-215 | 167-185 | 159.8-182 |
| Resolution (SASRES) (Å) | 59 ± 4 | 61 ± 4 | 45 ± 3 | 45 ± 3 |
| DAMMIN (Refinement of damstart.pdb) | | | | |
| q-range for fitting (Å$^{-1}$) | 0.00391-0.18185 | 0.00428-0.18669 | 0.00279-0.19258 | 0.00279-0.19587 |

| | | | | |
|---|---|---|---|---|
| Symmetry, anisotropy assumptions | P1, unknown | P1, unknown | P1, unknown | P1, unknown |
| Ambiguity score (AMBIMETER) | 0.4771 | 1.505 | 0.4771 | 0.4771 |
| $\chi^2$ | 1.43 | 4.359 | 2.716 | 7.365 |
| Model MW estimate (ratio to expected) | 271.9 (1.52) | 232.9 (1.30) | 230.4 (1.37) | 213.8 (1.27) |
| Model $R_g$ range | 45.2 | 46.36 | 42.62 | 42.29 |
| Model $D_{max}$ range | 183.5 | 173.5 | 167.2 | 159.8 |

1. K. Inamori, T. Yoshida-Moriguchi, Y. Hara, M. E. Anderson, L. Yu, K. P. Campbell, Dystroglycan Function Requires Xylosyl- and Glucuronyltransferase Activities of LARGE. *Science.* **335**, 93–96 (2012).

2. N. Kirby, N. Cowieson, A. M. Hawley, S. T. Mudie, D. J. McGillivray, M. Kusel, V. Samardzic-Boban, T. M. Ryan, Improved radiation dose efficiency in solution SAXS using a sheath flow sample environment. *Acta Crystallogr. Sect. D*. **72**, 1254–1266 (2016).

3. J. B. Hopkins, R. E. Gillilan, S. Skou, BioXTAS RAW: improvements to a free open-source program for small-angle X-ray scattering data reduction and analysis. *J. Appl. Crystallogr.* **50**, 1545–1553 (2017).

4. S. P. Meisburger, A. B. Taylor, C. A. Khan, S. Zhang, P. F. Fitzpatrick, N. Ando, Domain Movements upon Activation of Phenylalanine Hydroxylase Characterized by Crystallography and Chromatography-Coupled Small-Angle X-ray Scattering. *J. Am. Chem. Soc.* **138**, 6506–6516 (2016).

5. D. Svergun, A. Semenyuk, GNOM: Small-Angle Scattering Data Processing by Means of Regularization Technique. *Moscow, Russ. Inst. Crystallogr. Acad. Sci. Russ.* (1993).

13. M. Varadi, S. Anyango, M. Deshpande, S. Nair, C. Natassia, G. Yordanova, D. Yuan, O. Stroe, G. Wood, A. Laydon, A. Žídek, T. Green, K. Tunyasuvunakool, S. Petersen, J. Jumper, E. Clancy, R. Green, A. Vora, M. Lutfi, M. Figurnov, A. Cowie, N. Hobbs, P. Kohli, G. Kleywegt, E. Birney, D. Hassabis, S. Velankar, AlphaFold Protein Structure Database: massively expanding the structural coverage of protein-sequence space with

high-accuracy models. *Nucleic Acids Res.* **50**, D439–D444 (2022).
